# Social brain fingerprinting: unique functional connectomes underpin interpersonal behaviours in non-autistic and autistic adults

**DOI:** 10.64898/2026.10.20.760343

**Authors:** Augusto Ielo, Kristína Czekóová, Gianpaolo Antonio Basile, Alberto Cacciola, Milan Brázdil, Daniel Shaw, Enrico Amico

## Abstract

Human social interaction requires continuous adaptation while preserving stable individual patterns of brain organisation. This study examined whether functional connectome fingerprinting captures person-specific organisation during reciprocal interaction and whether its spatial architecture relates to social role and autistic traits. fMRI data from 70 adults forming 35 dyads were analysed during an iterated Ultimatum Game: 19 non-autistic Proposer–non-autistic Responder dyads and 16 non-autistic Proposer–autistic Responder dyads. Functional connectivity across 400 cortical parcels was used to assess whole-brain identifiability and edge-wise reliability. Connectomes remained identifiable across datasets, roles, and task conditions, with no reduction in global identifiability in mixed dyads. Edge-wise analyses revealed role-related and dataset-dependent differences across distributed large-scale networks. Role-sensitive connectivity features were associated with Autism-Spectrum Quotient scores, supported out-of-sample prediction, and topology-derived features classified AQ-threshold status above chance. Stable connectomic signatures therefore persist during reciprocal interaction while their network organisation varies with social role and autistic traits.

## Introduction

Human social interaction is not a sequence of isolated responses but a dynamic and non-linear process in which each individual continuously adjusts their behaviour in response to their interaction partner. Across repeated exchanges, actions are shaped not only by immediate goals but also by an ongoing evaluation of the partner’s prior behaviour, inferred intentions, and likely future responses^1,2^. In this sense, reciprocal interaction is inherently relational and temporally extended: each person’s behaviour is simultaneously a consequence of the other’s actions and a source of constraint for what follows next^3,4^. This mutual dependency is especially evident in dyadic contexts that require coordination, negotiation, or cooperation, where successful interaction depends on the capacity to update behaviour online as the exchange unfolds^2,4^. For social neuroscience, this implies that the mechanisms supporting interpersonal behaviour cannot be fully characterised from static or one-sided situations alone, because the core features of reciprocity emerge over time through active engagement between interacting individuals^3,5^.

Autism is defined behaviourally by differences in social communication and interaction. Among these characteristics, atypical social–emotional reciprocity is one of the most central and clinically informative features^6–9^. In practice, such differences are often expressed as reduced back-and-forth exchange, altered responsiveness to a partner’s behaviour, or less flexible adjustment of one’s actions within an ongoing interaction^6,10^. For this reason, reciprocity is not simply one social function among others, but a useful level at which to investigate how interpersonal difficulties unfold in real time^10,11^. Nonetheless, progress in identifying the mechanisms underlying these difficulties has been limited by the fact that many experimental paradigms isolate individuals from the reciprocal contexts in which social behaviour normally develops and is expressed^11,12^. As a result, atypical social behaviour in autism may appear less clearly in conventional laboratory settings than in naturalistic situations that require active, contingent exchange with another person^10,12^. A mechanistic account of autism-related social differences therefore requires approaches that treat reciprocity as an interactive process rather than as a static trait measured outside the dynamics of real-world exchange^10–12^.

The iterated Ultimatum Game (iUG) provides a controlled yet ecological valid interactive (“second-person”) paradigm for investigating reciprocity during ongoing social interaction^1,4^. In each exchange, the Proposer decides how to divide a sum of money between the two players, and the Responder decides whether to accept or reject the offer; acceptance implements the proposed division, whereas rejection leaves both players without any payoff^4^. Importantly, modifications of the Proposer’s choice set can introduce different monetary contexts that elicit variability in expressions of reciprocity. In the modified iUG, two such contexts are included. In Proposer–Responder (PR) exchanges, Proposers choose between advantageous and disadvantageous inequity (e.g., 60:40 vs 40:60), such that selecting the more generous option involves a greater personal cost and can therefore signal stronger cooperative intent. In Proposer–Proposer (PP) exchanges, both alternatives favour the Proposer but differ in the magnitude of advantageous inequity (e.g., 60:40 vs 70:30), such that selecting the more selfish option can indicate lower cooperative intent. Because both Proposers’ and Responders’ decisions depend partly on the alternatives available and on the inferred motivations of their partner, the PP and PR contexts are intended to elicit different expressions of positive and negative reciprocity across repeated exchanges. By repeating this exchange across multiple rounds, the iUG allows each player’s behaviour to depend not only on the current offer, but also on the history of previous interactions and expectations about the partner’s future behaviour^4^. In contrast to one-shot paradigms, this iterated structure allows reciprocity to emerge dynamically across rounds and makes it possible to distinguish the role-dependent contributions of Proposers and Responders^4,13^. The interactive structure of the iUG also makes it suitable for second-person neuroscience^14^; neural activity can be recorded from members of an interacting dyad^14^, allowing functional brain organisation to be examined in real-time while they engage in a controlled sequence of reciprocal exchanges^4,13,14^.Previous work using the iUG in brain imaging studies has shown that the choices of both Proposers and Responders across repeated economic exchanges can be estimated not only as an interactional process of mutual reciprocity rather than a series of one-off decisions, but also by the strength of localised functional brain connectivity^4,15^. More recently, this approach was extended to dyads comprising autistic and non-autistic players, showing that autistic Responders expressed reduced behavioural reciprocity and spent less time in states of dynamic brain connectivity characterised by flexible integration and segregation among large-scale control and default-mode networks^11,15^. Together, these findings demonstrate the value of the iUG and dual-fMRI hyperscanning paradigms for identifying neural mechanisms underpinning live social exchange, and suggest that autism-related social differences may be detectable in the organisation and temporal reconfiguration of large-scale functional brain networks during interaction^11,15^.

Although prior studies have begun to reveal the neural correlates of behavioural reciprocity, it remains unknown if reciprocal social exchange may be linked to reliable, individual-specific patterns of whole-brain functional organisation that can be quantified at the connectome level^14,15^. One way to study this is to investigate whether large-scale patterns of functional connectivity retain stable, individual-specific structures even while behaviour changes from moment to moment, as it does during social interaction. Work on functional connectome fingerprinting has shown that whole-brain connectivity profiles can identify individuals reliably across repeated measurements, indicating that the functional connectome contains reproducible signatures of person-specific brain organisation^16^. Importantly, this identifiability can be observed across multiple task contexts, suggesting that stable individual structure coexists with ongoing state-dependent reconfiguration^17,18^. Moreover, connectome fingerprinting is not only a measure of reliability or uniqueness; accumulating evidence indicates that the connections contributing most strongly to individual identifiability are behaviourally relevant and may offer a useful bridge between network architecture, cognitive abilities and clinical variations^19,20^. These developments make fingerprinting especially attractive for the study of social interaction, where a crucial yet unresolved question is whether the neural architecture of reciprocal exchange preserves stable individual signatures and, if so, whether the spatial organisation of those signatures differ as a function of interactional role (Proposer vs Responder) and autism-related traits^16,19,20^.

Whole-brain identifiability provides a useful summary of whether functional connectomes retain individual specificity, but it does not indicate which connections are responsible for that individuality or how those informative edges are organised across the connectome^16,17^. This is important because comparable levels of global identifiability can in principle arise from different underlying spatial configurations of reliable functional connections^17,20^. Recent work in connectome fingerprinting has therefore moved beyond whole-brain summary indices to examine the edge-wise spatial specificity of fingerprints, typically by quantifying the test– retest reliability of individual connections and identifying the subsets of edges that contribute most strongly to person-level distinctiveness. Such analyses make it possible to ask whether fingerprint architecture is concentrated within particular large-scale systems, redistributed across network interactions, or reorganised as a function of task demands or clinical status^18–20^. In the present context, this method is especially relevant because reciprocal social exchange may preserve overall connectome identifiability while altering the spatial distribution of the edges that are most stable in Proposers and Responders. An edge-wise approach is therefore necessary to determine whether role-related and autism-related differences emerge not at the level of global fingerprint strength, but in the network topography of temporally reliable functional connections^17,18,20^.

If role-related differences in edge-wise fingerprint architecture capture meaningful aspects of neural organisation during social interactions, they should not only distinguish interactional roles or datasets at the group level but also relate to inter-individual variation in autistic traits. This expectation is supported by a broader literature showing that person-specific features of functional connectomes are not merely reliable identifiers but can also carry behaviourally and clinically informative signal. In particular, fingerprint-derived architectures and closely related connectivity features have been linked to cognitive performance, developmental variation, and disease-related individual differences, suggesting that the same network properties that support identifiability may also encode meaningful variation in phenotype^17,19,20^. In this context, autistic traits provide a relevant continuous dimension because they vary within non-autistic and autistic individuals and may reflect how strongly interaction-related neural organisation is coupled to the social and cognitive demands of the task. This individual-level perspective is particularly relevant in autism, where functional brain organisation is markedly heterogeneous and population-level contrasts may obscure substantial inter-individual variability in connectivity patterns^21^. Notably, much of the evidence characterising such heterogeneity has been derived from resting-state functional connectivity, whereas task-based data provide the opportunity to examine individual differences in functional organisation under a common, experimentally constrained behavioural context. A second implication follows from this point: if role-sensitive connectivity features are related to trait variation, they may also support subject-level prediction and classification, moving the analysis from descriptive group contrasts to individual-level modelling. Progressing from the identification of stable network signatures to testing their predictive value has already proved informative in other clinical settings, where fingerprint-informed topological features have been used to predict clinically relevant outcomes^22–24^. Applied to the present study, this framework makes it possible to test whether the topology of these role-sensitive connectivity patterns contains information not only about Proposer and Responder organisation, but also about individual variation in autistic traits and whether these patterns classification according to psychometric-based thresholds.

Against this background, the present study examined whether functional connectome fingerprinting can capture stable and behaviourally meaningful signatures of brain organisation during live social interaction. Two independent datasets were analysed, both of which comprised behavioural and functional brain imaging data acquired with functional magnetic resonance imaging (fMRI) from players who interacted with one another on the iUG. One dataset was collected from dyads of non-autistic Proposers and non-autistic Responders (NP-NR), and the other comprised dyads of non-autistic Proposers and autistic Responders (NP-AR). First, whole-brain identifiability was assessed to determine whether individual functional connectomes remained distinguishable across task conditions and participant groups. Second, edge-wise spatial specificity was examined using intraclass correlation (ICC) to test whether the organisation of the most temporally reliable connections differed between Proposers and Responders within each dataset. Third, the resulting role-difference maps were compared between datasets to establish whether NP-NR and mixed NP-AR dyads showed distinct large-scale patterns of differential ICC. Since the individual-level analyses were designed to examine whether role-sensitive connectomic organisation captured stable variation in autism-related phenotype, rather than moment-to-moment variation in reciprocal behaviour, autistic traits were examined using AQ scores. Finally, focusing on the autistic sample, the study tested whether role-sensitive connectivity and topological features were associated with autistic traits and whether they supported individual-level prediction of continuous AQ scores and AQ-based group classification.

Together, these analyses were designed to determine whether reciprocal social exchange preserves identifiable whole-brain connectome structure, whether role-sensitive fingerprint architecture differs as a function of dataset, and whether these interaction-related connectomic patterns carry information relevant to autistic-traits variation.

## Results

Figure 1 summarises the main analysis workflow. Two fMRI datasets were acquired from independent samples of individuals while they interacted with another player on the iUG: one sample comprised 19 pairs of NP-NR, and the other comprising 16 pairs of NP-AR. For each of the 70 individuals, functional connectivity (FC) matrices among 400 cortical parcels were derived separately for each of three task conditions and used for whole-brain identifiability and edge-wise ICC analyses.

**Figure 1.**
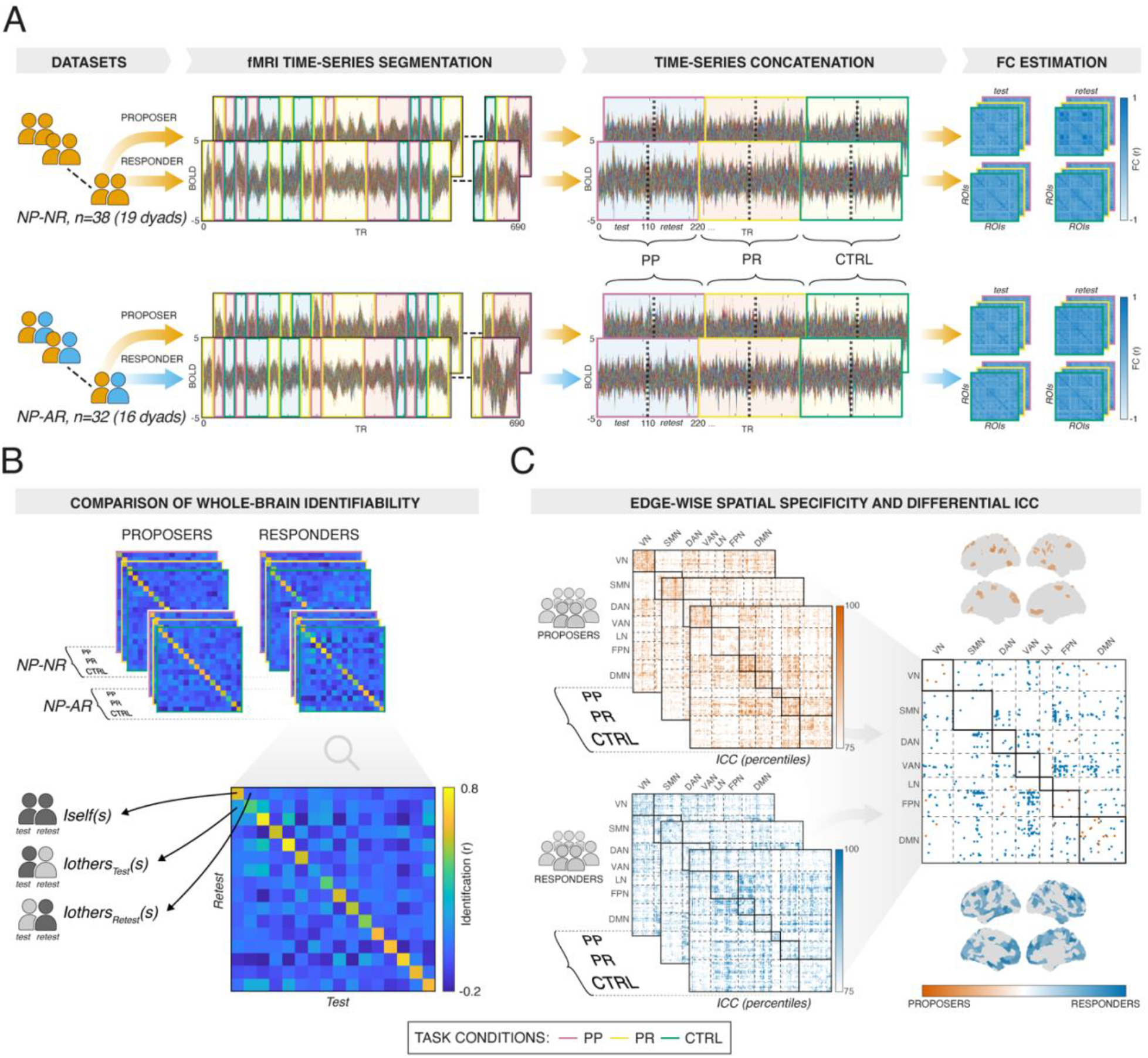
Study overview. A) Two fMRI datasets were analysed, one acquired from 19 dyads of non-autistic individuals (NP-NR) and the other from 16 dyads composed of a non-autistic Proposer and an autistic Responder (NP-AR). For each participant, fMRI time-series were extracted from 400 cortical parcels representing all non-overlapping nodes of seven intrinsic functional networks. These time-series were then segmented according to three task conditions (CTRL, PP, PR) and concatenated. Functional connectivity (FC) matrices were then estimated for each half of the condition-specific concatenated time-series, producing test and retest halves to enable identifiability analyses. B) Whole-brain identifiability was assessed separately for Proposers and Responders by computing correlations between FC matrices obtained from test and retest halves. The resulting identifiability matrices quantify whether an individual’s test FC pattern is more similar to their own retest FC (self-identifiability) compared to those of other participants (others-identifiability). Statistical evaluation of identifiability differences included Welch two-sample t-tests for between-group contrasts and paired t-tests for within-subject comparisons between task conditions. C) For the NP-NR dataset, edge-wise spatial specificity was quantified using the intraclass correlation coefficient (ICC) to estimate the stability of each functional connection across test and retest halves. A differential ICC analysis was then performed using a permutational null model to identify only those edges whose ICC was significantly higher in Proposers or Responders.Legend: NP-NR – non-autistic Proposer–non-autistic Responder; NP-AR – non-autistic Proposer–autistic Responder; BOLD – Blood oxygenation level dependent; TR – Repetition Time; CTRL – Control condition; PP – Proposer-Proposer condition; PR – Proposer-Responder condition; FC – Functional connectivity; ROI – Region of Interest; ICC – Intraclass correlation coefficient; VN – Visual Network; SMN – Somatomotor Network; DAN – Dorsal-attention Network; VAN – Ventral-attention Network; LN – Limbic Network; FPN – Fronto-parietal Network; DMN – Default Mode Network; .

### Whole-brain identifiability

Whole-brain identifiability matrices were examined separately for Proposers and Responders of NP-NR dyads, and Responders of NP-NR and NP-AR dyads in each task condition (Figure 2). Across all groups and task conditions, values along the main diagonal exceeded off-diagonal values, indicating higher similarity between each participant’s test and retest FC profile compared to those of other individuals.

**Figure 2.**
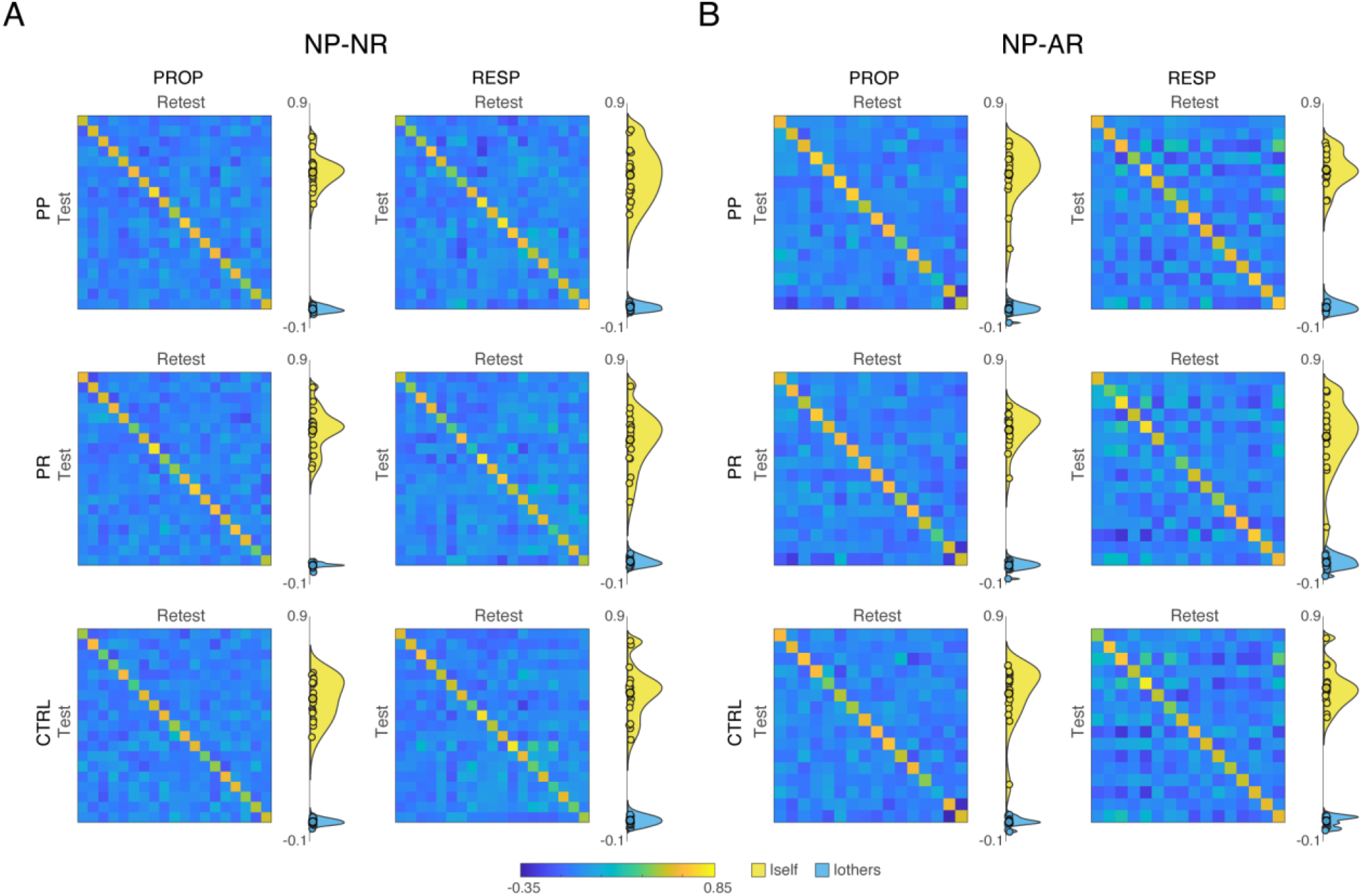
Whole-brain identifiability. Identifiability matrices are shown for the NP-NR dataset (A) and the NP-AR dataset (B), separately for proposers (PROP) and responders (RESP), and for each task condition (PP, PR, CTRL). In each matrix, rows correspond to test FC profiles and columns to retest FC profiles; warmer colours indicate higher similarity. For each matrix, the adjacent side raincloud plot shows the distributions of self-similarity (Iself; diagonal values) and others-similarity (Iothers; off-diagonal subject-level averages). Across datasets, roles and task conditions, diagonal values exceeded off-diagonal values, indicating higher similarity between each participant’s test and retest connectome than between that participant and other individuals. Colour scales are fixed across panels.

This pattern was also reflected in the summary identifiability indices. In the pooled residualised dataset used for the main identifiability analyses, mean Iself ranged from 0.54 to 0.59 in NP-NR Proposers, from 0.54 to 0.58 in NP-NR Responders, from 0.56 to 0.59 in NP-AR Proposers, and from 0.56 to 0.60 in NP-AR Responders. Mean Iothers values were close to zero across all groups and task conditions. Differential identifiability expressed as Idiff-norm was positive in all cases, ranging from 7.25 to 9.43 in NP-NR participants and from 5.32 to 7.77 in NP-AR participants. Identification success rate was 100% in all groups and conditions, meaning that, in every case, each participant’s test FC profile was matched most strongly to that participant’s own retest FC profile rather than to the retest profile of any other participant (Table 1).

**Table 1.** Whole-brain identifiability summary across groups and task conditions. Mean self-identifiability (Iself), mean others-identifiability (Iothers), differential identifiability expressed as percentage difference (Idiff), normalised differential identifiability (Idiff-norm), and identification success rate are shown for each group and task condition in the pooled residualised dataset used for the main identifiability analyses. Values for Iself and Iothers are reported as mean ± SD. Idiff-norm is reported with its 95% bootstrap confidence interval. In all cases, there was 100% success in identifiability.

| Dataset | Role | Condition | *I*self (mean ± SD) | *I*others (mean ± SD) | *I*diff (%) | *I*diff-norm (95% CI) |
| --- | --- | --- | --- | --- | --- | --- |
| NP-NR | PROP | CTRL | 0.54 ± 0.09 | -0.01 ± 0.01 | 52.03 | 7.78 (6.90–9.10) |
| NP-NR | PROP | PP | 0.59 ± 0.07 | -0.02 ± 0.01 | 57.66 | 9.43 (8.49–10.76) |
| NP-NR | PROP | PR | 0.59 ± 0.09 | -0.02 ± 0.01 | 57.10 | 8.93 (7.95–10.49) |
| NP-NR | RESP | CTRL | 0.56 ± 0.11 | -0.01 ± 0.02 | 54.35 | 7.25 (5.71–9.37) |
| NP-NR | RESP | PP | 0.58 ± 0.11 | -0.01 ± 0.01 | 56.82 | 7.92 (6.70–9.71) |
| NP-NR | RESP | PR | 0.54 ± 0.12 | 0.00 ± 0.02 | 52.99 | 7.05 (5.71–8.82) |
| NP-AR | PROP | CTRL | 0.56 ± 0.13 | -0.01 ± 0.02 | 53.89 | 7.11 (5.64–9.45) |
| NP-AR | PROP | PP | 0.59 ± 0.12 | -0.02 ± 0.02 | 56.74 | 7.59 (5.99–10.22) |
| NP-AR | PROP | PR | 0.59 ± 0.08 | -0.02 ± 0.02 | 57.00 | 7.77 (6.46–9.90) |
| NP-AR | RESP | CTRL | 0.58 ± 0.09 | -0.01 ± 0.02 | 56.26 | 5.60 (4.21–8.30) |
| NP-AR | RESP | PP | 0.60 ± 0.08 | -0.01 ± 0.02 | 58.84 | 5.89 (4.56–8.08) |
| NP-AR | RESP | PR | 0.56 ± 0.15 | 0.00 ± 0.03 | 53.82 | 5.32 (4.02–7.82) |

Across-condition comparisons within each group showed limited evidence of modulation by task condition; no across-condition contrast survived false discovery rate (FDR) correction for Iself. For Iothers, significant effects were observed only for Responders of NP-NR dyads, in whom Iothers was higher in the PR compared to both the CTRL and PP conditions (q = 0.001 for both contrasts; Supplementary Table S1). No other across-condition contrast survived FDR correction.

Between-dataset comparisons of identifiability components within role did not indicate differences between NP-NR and NP-AR datasets. Across the CTRL, PP and PR conditions, neither Iself nor Iothers differed significantly between Proposers or Responders of NP-NR and NP-AR dyads after FDR correction, and permutation tests showed the same pattern (Supplementary Table S2).

### Edge-wise spatial specificity: role-related ICC differences within each dataset

Edge-wise spatial specificity was assessed in the pooled residualised data by comparing ICC profiles between Proposers and Responders separately within the NP-NR and NP-AR datasets (Figure 3). In both datasets, significant role-related ICC differences were detected in all task conditions and in the condition-averaged map (AVG).

**Figure 3.**
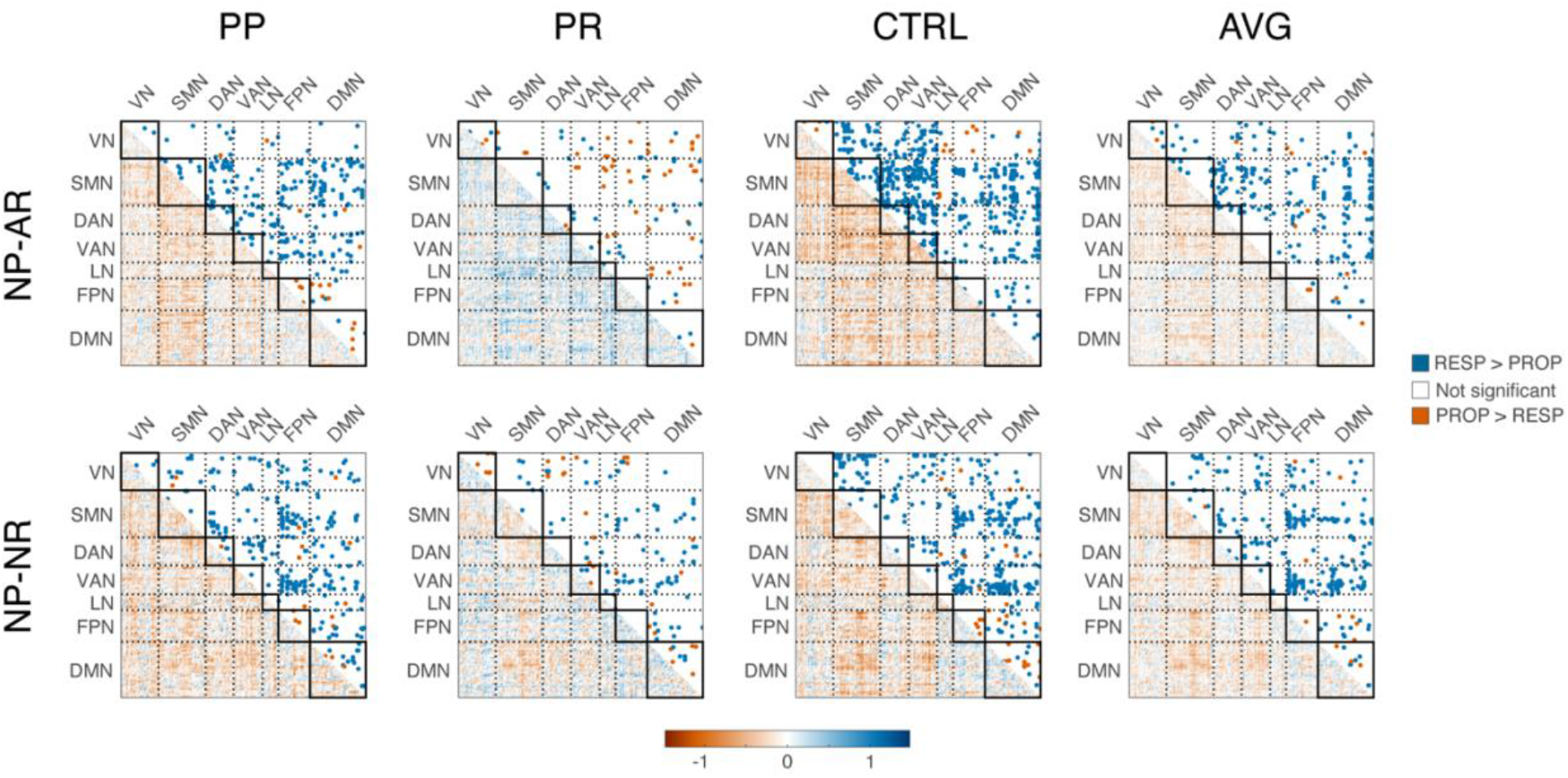
Within-dataset role-related differences in edge-wise ICC. Columns report results for each (CTRL, PP and PR), and the condition-averaged map (AVG). Rows report the two datasets (NP-NR and NP-AR). In each matrix, the lower triangle shows the differential edge-wise ICC map (ΔICC; PROP − RESP) and the upper triangle shows the corresponding significance mask after permutation testing with FDR correction; blue indicates edges with higher ICC in Responders compared with Proposers (RESP > PROP), orange indicates the reverse (PROP > RESP), and white indicates non-significant differences. Matrices are displayed in Yeo network order, with solid black squares marking within-network blocks and dotted lines marking network boundaries.

In the NP-NR dataset, 673 edges survived FDR correction in the CTRL condition, 281 in the PP condition, 98 in the PR condition, and 276 in the condition-averaged (AVG) map. In CTRL, PP, and AVG, most significant effects were in the Responders > Proposers direction (658, 263, and 263 edges, respectively). In PR, 46 edges showed RESP > PROP effects and 52 showed Proposers > Responders effects. In the NP-AR dataset, 431 edges survived correction in CTRL, 307 in PP, 134 in PR, and 307 in AVG. In CTRL, PP, and AVG, most significant effects were also in the Responders > Proposers direction (397, 289, and 294 edges, respectively), whereas fewer edges showed Proposers > Responders effects.

At the network level, the significant Proposer vs Responder ICC differences involved multiple large-scale systems (Supplementary Figure S1). In the NP-NR dataset, Responders > Proposers effects were observed in connections involving somatomotor, dorsal attention, ventral attention, and default mode systems, particularly in the CTRL and PP conditions. Proposers > Responders effects were fewer and observed mainly in the PR condition. In the NP-AR dataset, Responders > Proposers effects also involved multiple between-network pairs, with contributions from visual, somatomotor, ventral attention, frontoparietal, and default mode systems in CTRL and PP, and fewer effects in PR.

### Network-level comparison: condition-dependent differences between datasets

To test whether Proposer vs Responder differential ICC differed between datasets, the dataset-specific role-difference maps were compared at the network-pair level (Supplementary Figure S2). Significant between-dataset effects differed across conditions: In CTRL, significant effects were observed only in the NP-NR > NP-AR direction and were located mainly in interactions involving limbic, frontoparietal, ventral attention, somatomotor, and default mode networks. In PP and PR, significant effects were observed only in the NP-AR > NP-NR direction; in PP, these involved interactions centred mainly on visual, somatomotor, dorsal attention, ventral attention, and frontoparietal networks, whereas in PR they were more restricted and involved a smaller set of pairs centred mainly on visual, dorsal attention, and frontoparietal interactions.

### Differential ICC patterns within the NP-AR dataset

In the dataset comprising pairs of non-autistic Proposers (NP) and autistic Responders (AR), analysed in the residualisation set B, significant Proposer vs Responder differences in edge-wise ICC were observed in all task conditions and in the condition-averaged map (AVG). In all four cases, the predominant direction was AR > NP, with 493, 305, 93, and 311 significant edges, respectively, compared with 34, 20, 32, and 22 in the NP > AR direction. At the network level, AR > NP effects in CTRL were mainly distributed across network interactions involving ventral attention, default mode, somatomotor, frontoparietal, and visual systems, with the strongest contributions in ventral attention–default mode, somatomotor–default mode, and somatomotor–frontoparietal interactions. In PP, the dominant AR > NP pattern again involved ventral attention, frontoparietal, default mode, and somatomotor systems, with prominent contributions from ventral attention–frontoparietal, somatomotor–frontoparietal, and ventral attention–default mode interactions. In PR, the overall number of significant edges was lower, but AR > NP effects remained more frequent than NP > AR, with contributions involving ventral attention, frontoparietal, default mode, limbic, and somatomotor systems. In AVG, the dominant pattern was again AR > NP, with strongest contributions from ventral attention– frontoparietal, somatomotor–frontoparietal, ventral attention–default mode, and somatomotor–default mode interactions (Figure 4).

**Figure 4.**
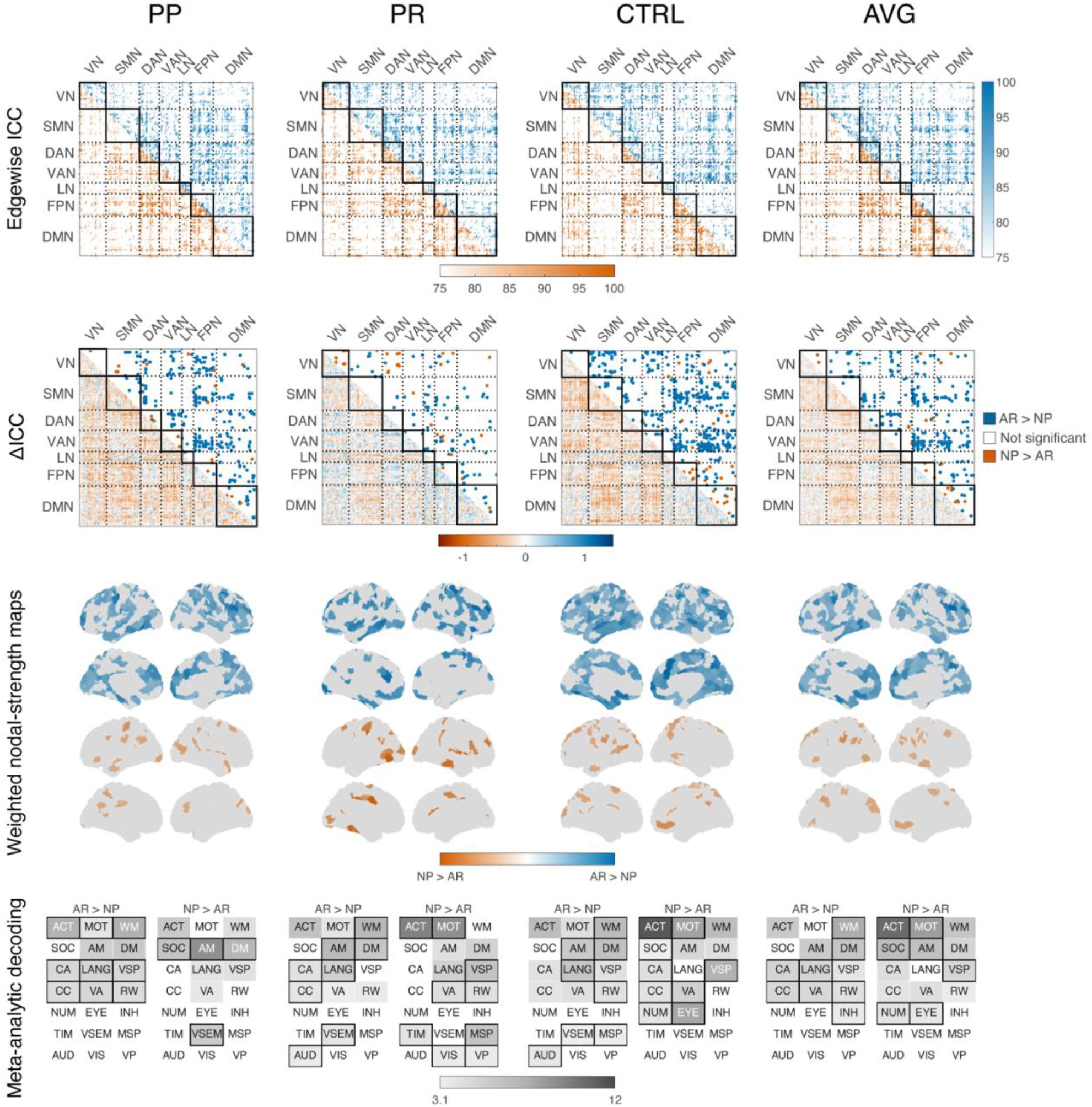
Differential ICC patterns within the NP-AR dataset. For each condition (CTRL, PP, PR) and the condition-averaged map (AVG), the figure shows four levels of representation of Proposer vs Responder ICC differences in the NP-AR dataset. In the first row, edge-wise ICC percentile maps are shown separately for non-autistic Proposers (NP) and autistic Responders (AR) within the same matrix; the lower triangle representing NP and the upper triangle representing AR. In the second row, the lower triangle displays the differential edge-wise ICC map (ΔICC), whereas the upper triangle shows the corresponding FDR-corrected significance mask. In the third row, weighted nodal-strength maps summarise the spatial distribution of significant effects on the cortical surface, separately for the two directions, using joint within-condition normalisation. Blue indicates edges with significantly higher ICC in Responders compared with Proposers (AR > NP), orange indicates the reverse (NP > AR), and white indicates non-significant differences. In the fourth row, meta-analytic decoding of the AR > NP and NP > AR differential ICC masks is shown for each condition. Cell shading represents the Neurosynth decoding z-statistic, with values displayed for z ≥ 3.1; unshaded cells indicate values below this threshold. Black outlines identify, for each matched cognitive term and condition, the direction with the higher raw z-statistic. Cognitive terms are indicated by acronyms within each cell. Legend: VN, visual network; SMN, somatomotor network; DAN, dorsal attention network; VAN, ventral attention network; LN, limbic network; FPN, frontoparietal network; DMN, default mode network; ACT, action; MOT, motor; WM, working memory; SOC, social cognition; AM, autobiographical memory; DM, declarative memory; CA, cued attention; LANG, language; VSP, visuospatial; CC, cognitive control; VA, visual attention; RW, reading/writing; NUM, numerical cognition; EYE, eye movements; INH, inhibition; VP, visual perception; VSEM, verbal semantics; MSP, multisensory processing; VIS, visual semantics; AUD, auditory processing; TIM, timing.

**Figure 5.**
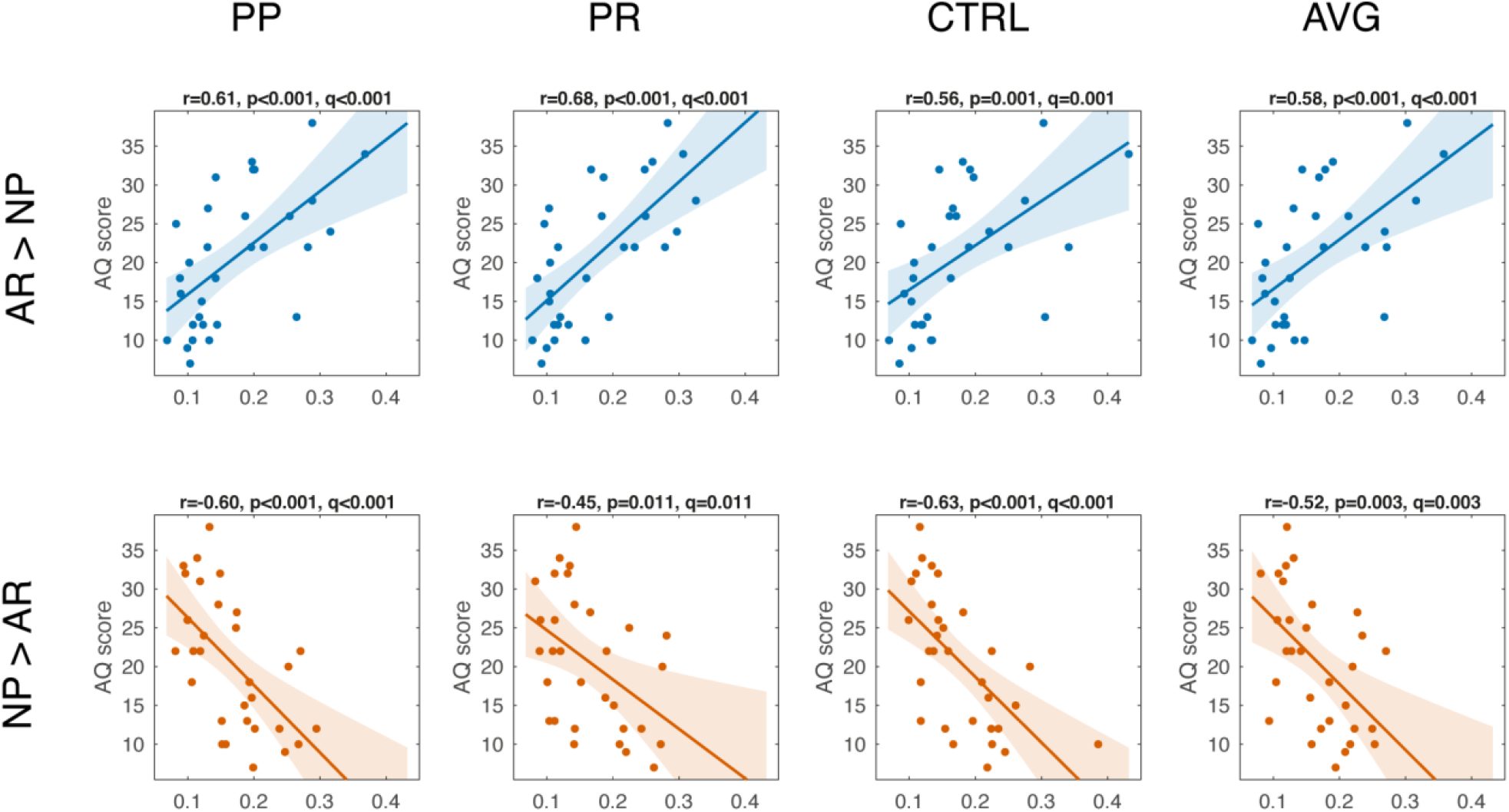
Associations between AQ scores and differential ICC-derived masked FC features in the NP-AR dataset. Columns report results for CTRL, PP, PR, and the condition-averaged map (AVG). Across non-autistic Proposers (NP) and autistic Responders (AR), rows show associations between AQ scores and masks in which edge-wise ICC is higher in AR relative to NP (top) **or those** expressing the opposing difference (NP > AR; bottom). For each panel, x-axes represent subject-level mean absolute FC within the corresponding common differential ICC mask, and y-axes represents AQ scores. Solid lines indicate linear fits and shaded areas indicate 95% confidence intervals. Pearson’s r, nominal p, and FDR-corrected q values are reported in each panel. Positive associations were observed for AR > NP masks, whereas negative associations were observed for NP > AR masks across all task conditions and AVG.

Meta-analytic decoding of the differential ICC masks for the NP-AR dataset indicated distinct but partially overlapping functional profiles for the two directions of effect. Across conditions, the AR > NP masks showed consistent decoding for terms related to action, motor, working memory, autobiographical memory, declarative memory, cued attention, language, visuospatial, cognitive control, visual attention, and reading/writing. Within this set, working memory, autobiographical memory, and declarative memory were represented across all four condition definitions, whereas other terms showed more condition-dependent variation. By contrast, the NP > AR masks also showed decoding for action, motor, working memory, autobiographical memory, declarative memory and social cognition, with the latter represented in CTRL, PP, and AVG. Additional condition-specific contributions included eye movements, visual perception, verbal semantics, multisensory processing, and timing.

### Autistic-traits analyses based on differential ICC-derived connectivity features

In Proposer vs Responder analyses for the NP-AR dataset, participant-level masked FC summaries derived from the common differential ICC masks across all task conditions and the condition-averaged (AVG) map were significantly associated with AQ scores across non-autistic Proposers and autistic Responders. For the AR > NP masks, AQ showed positive correlations in CTRL (r = 0.56, q < 0.001), PP (r = 0.61, q < 0.001), PR (r = 0.68, q < 0.001), and AVG (r = 0.59, q < 0.001). For the NP > AR masks, AQ showed negative correlations in CTRL (r = −0.63, q < 0.001), PP (r = −0.60, q < 0.001), PR (r = −0.45, q = 0.011), and AVG (r = −0.52, q = 0.003). Thus, higher AQ scores were associated with greater mean absolute FC within masks showing higher ICC in autistic Responders compared with non-autistic Proposers, and with lower mean absolute FC within masks showing higher ICC in Proposers than Responders.

### Prediction of AQ from differential ICC-derived connectivity features

To test whether differential ICC-derived masked connectivity features supported out-of-sample prediction of autistic traits, we next fitted leave-one-out linear models in the Set B dataset using fold-specific masks (Figure 6). Significant associations between observed and predicted AQ scores were detected for all task conditions and the condition-averaged map (AVG), in both mask directions. For the AR > NP masks, observed-versus-predicted AQ correlations were significant in CTRL (r = 0.43, R^2^ = 0.18, q = 0.019), PP (r = 0.54, R^2^ = 0.30, q = 0.003), PR (r = 0.64, R^2^ = 0.41, q = 0.001), and AVG (r = 0.49, R^2^ = 0.24, q = 0.008). For the NP > AR masks, significant correlations were also observed in CTRL (r = 0.60, R^2^ = 0.36, q = 0.001), PP (r = 0.54, R^2^ = 0.29, q = 0.003), PR (r = 0.40, R^2^ = 0.16, q = 0.025), and AVG (r = 0.43, R^2^ = 0.18, q = 0.019). Relative to a baseline model predicting the sample mean AQ score, all leave-one-out models showed lower RMSE and MAE, indicating that fold-specific differential ICC masks retained predictive information beyond a trivial mean-based prediction.

**Figure 6.**
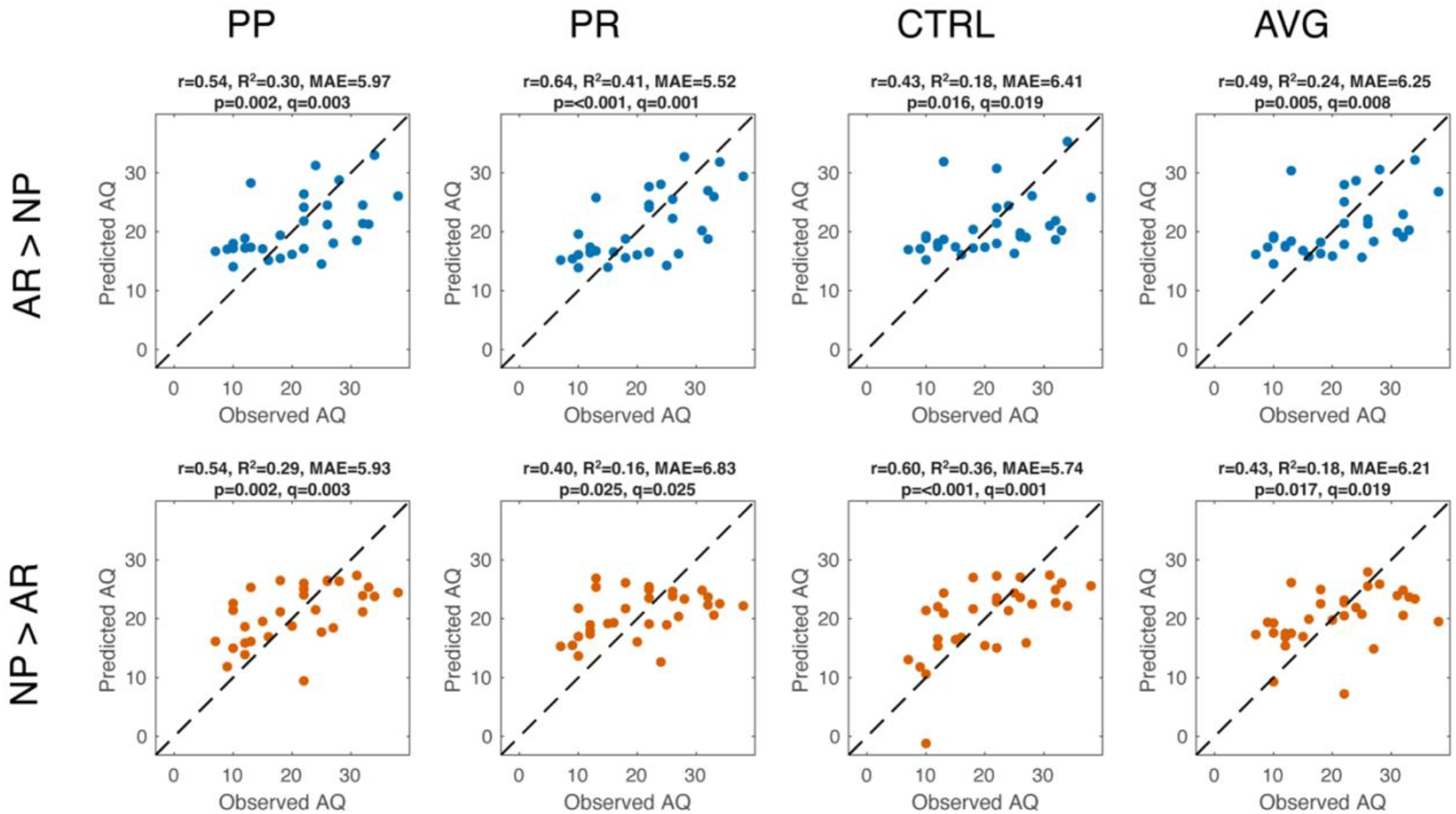
Leave-one-out prediction of AQ scores from differential ICC-derived masked FC features in the NP-AR dataset. Columns report results for CTRL, PP, PR, and the condition-averaged map (AVG). The top row shows leave-one-out prediction results for masks in which edge-wise ICC was higher in responders than in proposers (RESP > PROP), whereas the bottom row shows results for masks in which edge-wise ICC was higher in proposers than in responders (PROP > RESP). In each panel, the x-axis represents observed AQ score and the y-axis represents the corresponding leave-one-out predicted AQ score. The dashed diagonal indicates the identity line. Pearson’s r, coefficient of determination (R^2^), mean absolute error (MAE), nominal p, and FDR-corrected q values are reported in each panel. Significant out-of-sample prediction was observed across all task conditions and in the condition-averaged map for both mask directions.

### Binary classification of AQ status from differential ICC-derived topology features

A further analysis examined whether AQ-threshold status (AQ ≥ 26 vs < 26) could be classified from differential ICC-derived topology features in autistc Responders (Figure 7). Across the six tested FC condition and mask-direction combinations, the best classification performance was obtained using PR functional connectivity within the condition-averaged RESP > PROP differential ICC mask. The linear SVM achieved an accuracy of 0.87, balanced accuracy of 0.89, AUC of 0.85, F1-score of 0.88, and MCC of 0.76. The corresponding confusion matrix (Figure 7A) shows that all individuals in the lower-AQ group were correctly classified (specificity = 1.00, 6/6), whereas 7 of 9 individuals in the higher-AQ group were correctly classified (sensitivity = 0.78). The ROC curve for this model is shown in Figure 7B. Permutation testing indicated that the observed AUC exceeded the empirical null distribution (p = 0.021; Figure 7C). By contrast, the remaining condition/mask combinations showed lower performance, with AUC values ranging from 0.33 to 0.79 and balanced accuracy values ranging from 0.31 to 0.60, indicating that discriminative information was concentrated in the PR connectivity pattern masked by the AR > NP average differential ICC topology.

**Figure 7.**
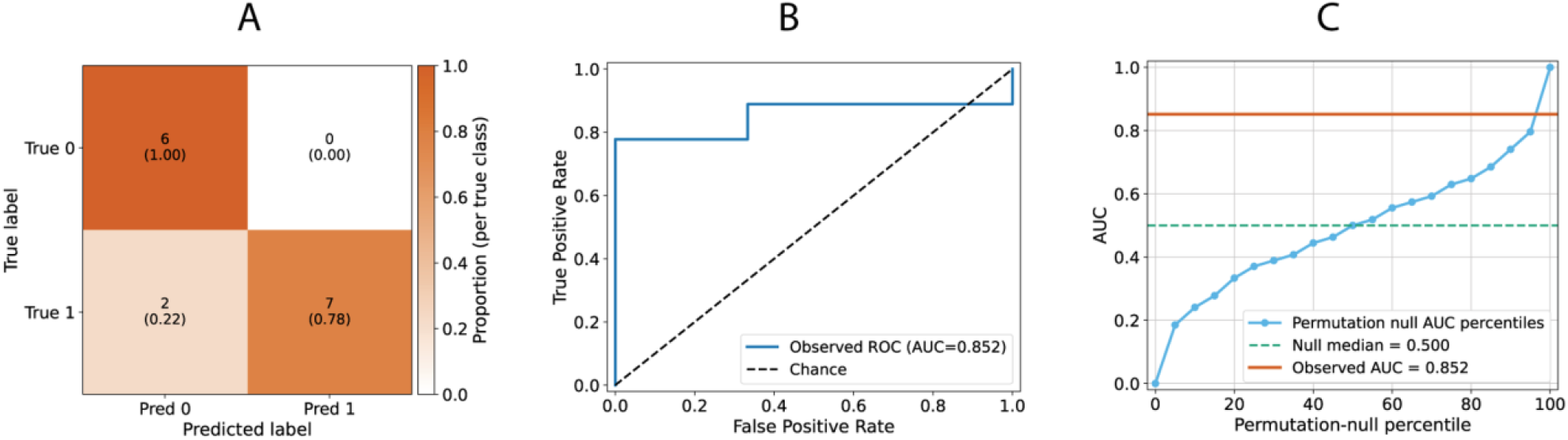
Binary classification of AQ status from differential ICC-derived topology features in NP-AR responders. Summary of the best-performing classifier identified across the tested conditions. Topology features were extracted from PR functional-connectivity matrices within the RESP > PROP condition-averaged differential ICC mask and used to classify participants according to AQ-threshold status (AQ ≥ 26 vs < 26) using a linear support vector machine in leave-one-out cross-validation. A) confusion matrix of outer-loop predictions, showing correct classification of all participants in the lower-AQ group (6/6; specificity = 1.00) and 7 of 9 participants in the higher-AQ group (sensitivity = 0.78). Cell colours represent within-class proportions. B) receiver operating characteristic curve for the same model, with an observed AUC = 0.852; the dashed diagonal indicates chance-level performance. C) permutation-test summary showing the observed AUC relative to the empirical null distribution obtained by label shuffling.

A threshold sensitivity analysis was performed for the best-performing classifier using alternative AQ cutoffs from 23 to 29 (Supplementary Table S3). Classification performance remained comparable for cut-offs between AQ ≥23 and AQ ≥26, with balanced accuracy of 0.89–0.90, AUC of 0.85–0.88, and significant permutation tests (p_perm_ = 0.012–0.021). Performance decreased at higher thresholds: AQ ≥27/28 yielded balanced accuracy = 0.53 and AUC = 0.63 (p_perm_ = 0.243), whereas AQ ≥29 yielded balanced accuracy = 0.67 and AUC = 0.78 (p_perm_ = 0.066). Thus, classification was robust across nearby lower thresholds but showed sensitivity to class composition at higher AQ cutoffs.

## Discussion

Brain fingerprinting was developed originally to determine whether individuals possess unique and stable patterns of whole-brain functional connectivity (FC) that can be reliably identified across repeated measurements^17^. Described as a new frontier in neuroscience, past studies have demonstrated that these connectomic signatures remain stable over prolonged periods of time^16,25,26^. Subsequent efforts have been focused on establishing the behavioural relevance of these individual differences by associating them with cognitive performance and behavioural phenotypes^27–26^. Despite these rapid advances, the relevance of connectome fingerprints for understanding real-world social behaviour remains largely unexplored.

To address this gap, the present study is the first to apply functional connectome fingerprinting to brain imaging data acquired during a second-person paradigm that emulates core characteristics of naturalistic social interaction. By introducing social brain fingerprinting as a novel framework, we establish its feasibility by making four principal contributions to the field of connectomics, computational and social neuroscience, and autism research. First, we demonstrate that functional connectome fingerprints remain highly identifiable during the continuous mutual adaptation of two individuals in real time. Second, we move beyond conventional fingerprinting by combining whole-brain identifiability analyses and edge-wise reliability mapping^17^. This combination revealed robust and reproducible role-dependent differences in individual signatures between Proposers and Responders, while leaving global fingerprint strength largely unchanged. Third, autism-related differences in functional brain connectivity were expressed primarily through the organisation of stable network interactions rather than through reduced connectome identifiability. Finally, fingerprinting-informed connectomic features were strongly associated with autistic traits, significantly predicted out-of-sample AQ scores, and enabled classification of AQ status.

### Stable individual connectomes during social interaction

A central question in social neuroscience concerns how stable neural organisation supports the dynamic processes that emerge during interpersonal interaction. Beyond establishing their uniqueness and temporal stability, connectome fingerprints have been shown to generalise across diverse cognitive domains including attention, working memory, language processing, and fluid intelligence, suggesting that person-specific functional organisation represents a stable and intrinsic feature of brain architecture^16,25,26^. Importantly, these signatures are not distributed uniformly across the connectome, with connections contributing most strongly to individual identification consistently concentrated within higher-order association systems, particularly the frontoparietal, default-mode, and attention networks^16,23,27^. Increasing interest in the real-world relevance of connectomic fingerprints, together with recognition of the limited behavioural utility of resting-state FC alone, has motivated a shift towards structured tasks and naturalistic paradigms that better capture behaviourally meaningful individual differences and improve behavioural prediction^30^. Recent evidence further indicates that dominant resting-state coactivation patterns can obscure connectivity features relevant to individual variability, while combining resting state with task-based FC enhances the reliability and behavioural relevance of individual connectivity estimates^31,32^.

Social interaction represents a particularly rigorous test of individual specificity because successful interaction requires ongoing adaptation to another person while simultaneously engaging neural systems central to real-world social functioning. Such adaptation is frequently associated with inter-brain alignment and neural coupling in interacting individuals^4^, a phenomenon that potentially implies a reduction in individual connectomic identifiability. In contrast, connectome fingerprints remained highly identifiable during our interactive iUG paradigm irrespective of role and interaction context, indicating that stable individual-specific patterns of functional organisation are preserved during reciprocal social exchange. These findings suggest that brain-to-brain coupling emerges from the encounter between two stable connectomic architectures^2^. Social interaction therefore appears to emerge through the flexible recruitment of stable person-specific connectomic architectures that act as scaffolds for interaction-related computations, extending previous proposals that social behaviour unfolds within the constraints imposed by highly stable large-scale functional architectures^33,34^.

### Role-dependent architecture of social interaction

A second major contribution of the present study is the demonstration that connectomic individuality is not fixed but can be expressed through distinct network configurations depending on social role. More broadly, the relevance of connectome fingerprinting lies not only in participant identification but also in determining whether individual signatures are behaviourally meaningful and predictive of cognitive or trait phenotypes. Although early work suggested some overlap between these dimensions, with the most discriminative networks also contributing to the prediction of abilities such as fluid intelligence^16^, more recent evidence suggests that the functional systems supporting participant identification and behavioural prediction are not necessarily the same^19^, and that behavioural prediction may benefit additionally from information derived from structure-function coupling^35^. Thus, high connectomic individuality is unlikely to be reducible to a single neural signature.

Traditional fingerprinting studies have focused predominantly on whole-brain identification accuracy as a global measure of individuality^16,17^. While highly informative, such approaches provide limited insight into where within the connectome individual signatures emerge. By combining identifiability analyses with edge-wise reliability mapping, we characterised not only the strength but also the spatial architecture of connectomic fingerprints. This approach proved particularly informative: although global fingerprint strength was comparable between Proposers and Responders, the network interactions contributing to that identifiability differed substantially across roles. In other words, participants remained equally identifiable irrespective of role, yet the connections supporting that identifiability were distributed differently across the connectome.

Alongside role-specific differences, several connectivity motifs were observed consistently across interaction conditions, particularly involving somatomotor, ventral attention and default mode networks. These networks support action planning and execution, processing of salient information, and self-referential and social-cognitive functions, respectively^36–38^. Their recurrent involvement suggests that stable individual differences during social interaction emerge from systems integrating perception, action, attention, and social evaluation.

Notably, this network profile closely resembles the architecture of connectomic individuality observed in non-social contexts. Fingerprint-defining connections in the general population are located most consistently within higher-order association systems, particularly the frontoparietal, default mode, dorsal and ventral attention networks^16,23,27^. The persistence of these systems during reciprocal interaction suggests that the neural basis of individuality remains stable across behavioural contexts, even when expressed through role-specific network configurations. These findings would have remained hidden under conventional fingerprinting approaches that focus solely on whole-brain identification metrics and demonstrate the value of examining the spatial architecture of connectomic fingerprints.

The predominance of Responder-related effects provides further insight into the neural mechanisms supporting social decision-making. Within the Ultimatum Game, Responders must evaluate fairness of the offer from the perspectives of both players while regulating their (negative) emotional response, infer Proposeŕs intentions, integrate information across previous interactions, and anticipate future consequences to select an appropriate behavioural response. Such demands recruit executive-control, attentional, mnemonic and self-referential systems^37,38^. Consistent with this interpretation, Responder-dominant reliability patterns prominently involved frontoparietal, ventralattention, default mode and somatomotor networks – systems linked by Neurosynth to working memory, autobiographical memory, cognitive control, action selection, and visuospatial processing. Taken together, these findings point toward connectomic individuality as a stable property of the brain that is flexibly reconfigured according to social role demands during interaction.

### Autism-related network organisation and clinical relevance

Another major implication of this work concerns the nature of autism-related differences in large-scale brain organisation. Research investigating functional connectivity in autism has produced highly heterogeneous results, with both hypo- and hyper-connectivity reported across social, attentional and executive-control systems^39–41^. These inconsistencies have been attributed to methodological variability, developmental factors or differences in sample composition. Increasingly, however, they are being interpreted in light of substantial within-group heterogeneity and functional idiosyncrasy, motivating a shift from conceptualising autism as a condition characterised by a shared connectivity phenotype towards a perspective acknowledging diverse and individually variable patterns of neural organisation^8,21,28,42,43^. Consistent with this perspective, participants with autism in the present study remained equally identifiable despite substantial reorganisation of the connections contributing to their fingerprints. Unlike conditions associated with reduced connectome identifiability^44–46^, this pattern more closely resembles reports of preserved fingerprint strength accompanied by reorganisation of fingerprint-defining connections found elsewhere^20^. This finding therefore suggests that atypical social cognition in autism does not arise from a global loss of connectomic individuality, but from alternative configurations of the network interactions that support it.

Interestingly, the identified alterations involve defaultmode, frontoparietal, dorsal and ventral attention, limbic, visual, and somatomotor networks. Several of these large-scale systems overlap with those identified as the most idiosyncratic in autism (DMN, SMN, VAN, DAN)^42,47^, as well as with those that contribute most strongly to highly individualised connectome fingerprints in the typically developing population^16,17,23^. These findings therefore suggest that autism may involve alternative configurations of the neural systems that ordinarily support individuality, but not a reduction of individuality.

Our final significant contribution builds on current efforts to enhance brain-behaviour prediction through behaviourally relevant imaging contexts^27,29–31^. Emerging evidence suggests that prediction of autistic traits can be improved when connectivity is measured during engaging tasks instead of passive conditions, such as unconstrained rest or free-viewing social attention paradigms^26,28^. However, evidence for the predictive advantage of behaviourally relevant contexts remains relatively scarce for social and autism-related phenotypes and these approaches have not yet been applied to reciprocal social interaction. Our findings extend this line of research toward stronger engagement by showing that individualised patterns of functional organisation expressed during reciprocal social exchanges are sufficiently informative to capture clinically relevant variation in autistic traits. Significant prediction of continuous AQ scores and above-chance classification of AQ-threshold status are in line with the notion that role-sensitive connectomic features contain behaviourally relevant information over and above participant identification.

Together, these findings provide compelling evidence that ecologically valid social interaction offers a uniquely informative context for revealing clinically and behaviourally relevant aspects of individual brain organisation that are not readily captured in conventional imaging paradigms.

### Limitations

An important limitation of this work is that the sample consisted exclusively of males. Although this strategy reduced phenotypic heterogeneity and increased group comparability, it prevents the generalisation of our findings to autistic females, who often present with distinct behavioural profiles (e.g., more frequent social camouflaging) and remain underdiagnosed until later in life^48,49^. Moreover, a substantial literature supports sex-related differences in FC and network organisation in autism^50^. Future studies should determine if and how the fingerprinting signatures and network-level alterations identified here generalise to females.

Although the combination of whole-brain identifiability analyses and edge-wise reliability mapping provides convergent evidence for the robustness of the observed effects, our sample size was relatively modest. Participant identification remained highly reliable, but AQ-status classification was more sensitive to sample composition—it remained stable across nearby lower AQ thresholds but decreased when higher thresholds altered class membership, suggesting sensitivity to the placement of individual participants around the classification threshold. Larger samples and independent validation are necessary before we can consider any clinical applications of our findings. Nevertheless, our findings contribute to broader efforts in precision psychiatry that seek to identify biologically informed markers capable of predicting individual differences rather than simply comparing diagnostic groups.

Third, within the NP-AR dataset, autistic participants were restricted to the Responder role whereas Proposers were always non-autistic. This might have introduced a mismatch between the socio-cognitive processing styles of the two interaction partners and potentially influenced their interaction dynamics^51^. Although comparisons with the NP-NR dataset allowed role-related effects to be contextualised, a fully factorial design including autistic participants in both interaction roles would provide a stronger separation of autism-related and role-related contributions. The absence of dyads consisting of two autistic interactants precluded evaluation of the observed interaction patterns in terms of autism-related characteristics per se in contrast to the reciprocal difficulties that can arise in mixed autistic-non-autistic dyads^52^.

Finally, the present findings were obtained within a single interactive paradigm and using previously acquired datasets. Replication across different forms of real-time social interaction will be important for establishing a broader relevance of the identified signatures.

## Methods

The present study reports secondary analyses of two previously acquired and published fMRI datasets. The NP-NR dataset was originally described in Shaw et al., 2018, whereas the NP-AR dataset was subsequently reported in Czekóová et al., 2025. The present study addresses a distinct research question by applying functional connectome fingerprinting analyses to these data. Full details of participant recruitment, the experimental protocol, and MRI acquisition are provided in the original reports^4,11^.

### Participants

The study included two dual-fMRI datasets acquired during the iterated Ultimatum Game (iUG). The first dataset, referred to hereafter as NP-NR, comprised 19 dyads of non-autistic participants^4^. The final NP-NR sample included 38 male participants forming 19 dyads after exclusion of one dyad for excessive head motion. Participants had a mean age of 24.6 years (SD = 3.7; range = 19.8–38.0), reported normal or corrected-to-normal vision, and had no history of neurological or psychiatric disorder.

The second dataset, hereafter referred to as the NP-AR dataset, comprised 16 dyads including a non-autistic Proposer and autistic Responder. The autistic Responders had a mean age of 24.5 years (SD = 6.1; range = 19.2–45.1). Data in the NP-AR dataset were acquired at the same institution as the NP-NR dataset and followed the same experimental procedure, MRI acquisition protocol, and preprocessing framework unless otherwise specified.

Written informed consent was obtained from all participants prior to scanning. Each of the studies in which these datasets were acquired was approved by the Ethics Committee of University Hospital Brno and/or the Research Ethics Committee of Masaryk University.

### Experimental task and procedure

Pairs of participants performed multiple rounds of an iUG with one another, throughout which one member of each dyad acted as Proposer and the other as Responder. In each round, the Proposer selected one of two possible monetary divisions and the Responder then chose whether to accept or reject the offer. If the Responder accepted the proposed division, the money was divided accordingly; if the Responder rejected it, neither player received any money. Three task conditions were analysed: a Proposer-Proposer condition (PP), a Proposer-Responder condition (PR), and a control condition (CTRL). In the PP condition, the two possible monetary divisions favoured the Proposer over the Responder (e.g., 60:40 | 70:30). In the PR condition, the Proposer instead chose between allocations involving advantageous or disadvantageous inequity for themself (e.g., 60:40 | 40:60). In the CTRL condition, participants made analogous choices involving colours rather than monetary allocations, and the Responder’s decision had no monetary consequences.

For both datasets, pairs of participants were introduced to one another for the first time on the day of scanning and remained in fixed Proposer and Responder roles throughout the session. The experiment consisted of two functional runs acquired within a single scanning session. Across the two runs, participants completed 120 iUG rounds (30 PP and 30 PR) and 60 control rounds. Each round consisted of a 4-s choice period, a 4-s offer period, and a 4-s decision period, followed by a jittered inter-trial interval of 2–4 s.

In the NP-AR dataset, both non-autistic and autistic participants completed the Autism Spectrum Quotient (AQ), a self-report questionnaire designed to quantify autistic traits^66^.

### MRI acquisition

MRI data were acquired at Masaryk University using two 3T Siemens Prisma scanners located in the same building (MRI-A and -B). In the NP-NR dataset, the two scanners were used in a counterbalanced manner across Proposer and Responder roles (i.e. 10 of the Responders were scanned in MRI-A and 9 were scanned in MRI-B). In the NP-AR dataset, all Responders were scanned on MRI-A and all Proposers on MRI-B. The scanners were identical in model, installation period, head-coil configuration and software environment, and used the exact same acquisition sequences and parameter settings. The scanners differed only in minor hardware characteristics that were not expected to introduce systematic differences beyond ordinary between-session variability. Potential scanner-related effects were explicitly controlled for by regressing out scanner identity from the functional connectivity data prior to subsequent analyses.

Functional images were acquired using a T2*-weighted echo-planar imaging sequence with a GRAPPA acceleration factor of 2, TR/TE = 2000/35 ms, and 690 volumes per run. A high-resolution T1-weighted structural scan was also acquired for anatomical localisation and co-registration.

### Preprocessing

For both datasets, fMRI data from each run was preprocessing following a pipeline that applied various tools packaged within FSL. This included motion correction performed with MCFLIRT. Residual motion- and physiology-related signal components were then identified automatically using SOCK (Bhaganagarapu et al. 2013, 2014) and subsequently regressed from the time series. Subsequent analyses treated scanner-related adjustment differently depending on the analytic set.

### Time-series segmentation and functional connectivity estimation

For each participant, whole-brain fMRI time series were segmented according to task condition (CTRL, PP, PR). All segments belonging to the same condition were then concatenated, producing condition-specific time series of 220 TRs (7.3 mins). Each condition-specific time series was then split into two equal halves of 110 TRs, which served as within-scan test and retest segments for connectome fingerprinting analyses. Functional connectivity (FC) matrices were estimated using a 400-region cortical parcellation based on the Schaefer atlas. For each parcel, the regional time series was obtained by averaging the preprocessed BOLD signal across all voxels within that parcel.

### Residualisation strategies

FC vectors were residualised in order to remove variance associated with nuisance covariates. Residualisation was performed separately for test and retest FC data within each task condition, using linear regression applied edge-wise across subjects. Here, test and retest corresponded to the first and second halves, respectively, of the same condition-specific time series derived from a single fMRI run. Three residualisation analysis sets (Set A, Set B, and Set C) were used, reflecting the structure of the datasets and the aims of the subsequent analyses.

In Set A, data from the NP-NR and NP-AR datasets were pooled and residualised with respect to intercept, participant role, scanner, and head motion. This set was used for analyses requiring direct comparison between datasets, because in the pooled sample scanner and role were not fully aliased. The regression model can be written as

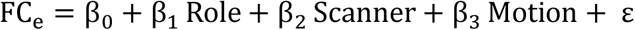

where FCe denotes the value of edge e across subjects, and the residual term ε was retained for subsequent analyses.

In Set B, only NP-AR participants were included, and FC vectors were residualised with respect to intercept and head motion:

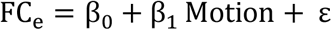

This set was used for NP-AR-only proposer-versus-responder analyses. In the NP-AR dataset, proposer and responder roles were tied to scanner assignment, such that all responders were scanned on one scanner and all proposers on the other. Under this design, scanner and role were perfectly confounded within the NP-AR-only sample. For this reason, scanner was not included as a covariate in Set B, so as to preserve the interpretable proposer-responder contrast.

In Set C, analyses of autistic traits were restricted to NP-AR responders only. For this subset, FC vectors were again residualised with respect to intercept and head motion:

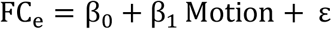

Although Sets B and C used the same covariate structure, they were estimated on different samples. Set B was fitted on the full NP-AR sample, whereas Set C was fitted on the NP-AR responder subset only.

### Connectome fingerprinting and identifiability

To quantify the subject-specific stability of functional connectomes, we applied a connectome fingerprinting framework based on pairwise similarity between vectorised functional connectivity (FC) profiles. For the main identifiability analyses reported here, FC vectors were taken from the pooled residualised dataset (Set A), in which edge-wise nuisance variance associated with intercept, participant role, scanner, and head motion had been removed prior to analysis. Identifiability was then assessed separately for each task condition (CTRL, PP, PR) and participant subgroup. Although identifiability analyses were also explored in the NP-AR-only motion-adjusted dataset, only the pooled Set A results are described in the main text.

For each participant, and separately for each condition, condition-specific FC matrices were obtained from the test and retest halves of the concatenated time series, Fisher z-transformed, and vectorised by retaining the upper triangular edges only. This provided one FC vector for the test half and one FC vector for the retest half for each participant and condition. As illustrated in the study workflow (Figure 1), identifiability analyses were performed separately for each subgroup, such that each identifiability matrix compared individuals within the same dataset-role combination.

For a given subgroup and condition, identifiability was assessed by constructing an N×N matrix M, where N is the number of participants in that subgroup and each entry represents the Pearson correlation between the test FC vector of one participant and the retest FC vector of another participant:

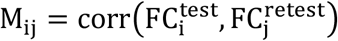

where FC_i_^test^ and FC_j_^retest^ denote the vectorised test and retest FC profiles of participants i and j, respectively. In this matrix, diagonal elements quantify within-subject similarity across the two halves of the scan, whereas off-diagonal elements quantify between-subject similarity. This formulation follows the standard identifiability framework based on all-against-all FC similarity.

For each participant s, self-identifiability (I_self_) was defined as the correlation between that participant’s own test and retest FC vectors:

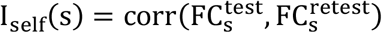

This metric quantifies the extent to which an individual’s connectome remains stable across the two halves of the same condition-specific time series.

Others-identifiability (Iothers) was defined, for each participant s, as the average similarity between that participant’s FC profile and the FC profiles of all other participants in the same identifiability matrix:

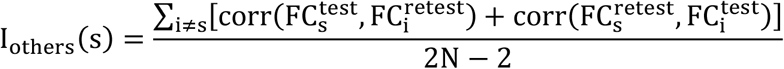

This quantity describes how similar a participant’s connectome is, on average, to the connectomes of the other individuals in the same subgroup. The symmetric formulation, averaging test-to-retest and retest-to-test matches, reduces directional bias due to the arbitrary assignment of halves as test and retest.
Differential identifiability (I_diff_) was then computed as the difference between the mean self-identifiability and the mean others-identifiability across participants:

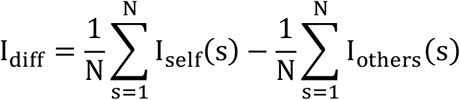

Higher values of I_diff_ indicate stronger separation between within-subject and between-subject FC similarity, and therefore greater connectome distinctiveness at the group level.

To obtain a scale-independent summary of identifiability, we also computed a normalised differential identifiability index (I_diff-norm_) as the standardised distance between the diagonal and off-diagonal distributions of the identifiability matrix:

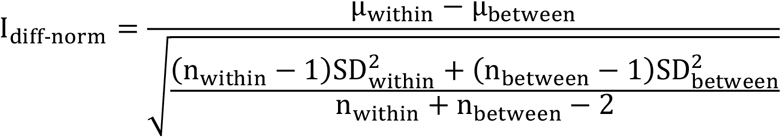

where μ_within_ and SD^2^within are the mean and variance of the diagonal entries of M, μ_between_ and SD^2^between are the mean and variance of the off-diagonal entries, and n_within_ and n_between_ are the corresponding numbers of observations. This normalised measure provides a scale-free effect size and facilitates comparison across conditions and participant subgroups.

Finally, we quantified identification success rate (SR) as the percentage of participants for whom the self-correlation exceeded all corresponding between-subject correlations, that is, the proportion of participants correctly matched to themselves across test and retest halves:

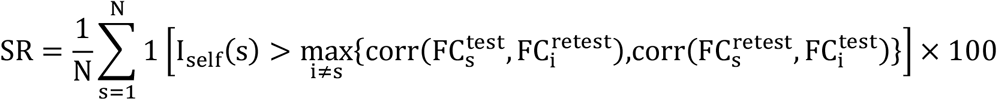

In the pooled Set A analysis, identifiability matrices were computed separately for NP-NR proposers, NP-NR responders, NP-AR proposers, and NP-AR responders within each condition. In addition to within-condition identifiability matrices, cross-condition test-versus-retest comparisons were also generated to summarise how fingerprint similarity varied across task conditions.

Statistical comparisons of identifiability components focused on Iself and Iothers. In the pooled analysis, between-dataset comparisons were performed separately within role and condition, comparing NP-NR proposers with NP-AR proposers and NP-NR responders with NP-AR responders. These contrasts were assessed using two-sided Welch two-sample t-tests, together with permutation tests based on label-shuffling of the group assignments and recomputation of the mean difference. In addition, within each subgroup, across-condition comparisons between CTRL, PP, and PR were assessed using two-sided paired t-tests and corresponding sign-flip permutation tests on paired differences. For these identifiability analyses, false discovery rate correction was applied separately to the families of Iself and Iothers tests.

### Edge-wise spatial specificity and differential ICC within datasets

To characterise the spatial specificity of connectome fingerprints, we quantified the temporal stability of each functional connection using the intraclass correlation coefficient (ICC). In this context, edge-wise ICC measures the extent to which the value of a given FC edge is more consistent within the same individual across test and retest than across different individuals. Following the standard formulation used in edge-wise fingerprinting studies, ICC was defined as

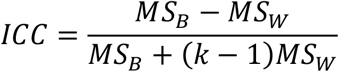

where MS_B_ is the between-subject mean square, MS_W_ is the within-subject mean square, and k is the number of repeated measurements. In the present study, k=2, corresponding to the test and retest halves of each condition-specific time series. Higher ICC values indicate that an edge shows greater between-subject variability relative to within-subject fluctuation and therefore contributes more strongly to subject-specific fingerprinting^20^.

ICC was stabilised by repeated random subsampling. For each condition and subgroup, 100 iterations were performed in which 80% of participants were sampled without replacement, edge-wise ICC was recomputed, and the resulting ICC profiles were retained. The final ICC estimate for a given edge was defined as the mean across these subsampling iterations. This procedure was applied separately to proposers and responders.

In the pooled residualised data, edge-wise spatial specificity was assessed separately within the NP-NR and NP-AR datasets. Thus, for each condition, four subgroup-specific ICC profiles were obtained: NP-NR proposers, NP-NR responders, NP-AR proposers, and NP-AR responders. Within each dataset, role-related differences in edge-wise spatial specificity were quantified as

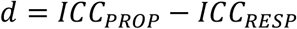

computed separately for CTRL, PP, and PR, as well as for an additional condition-averaged summary map obtained by averaging the three condition-specific contrasts. Positive values of d indicate greater ICC in proposers than in responders, whereas negative values indicate greater ICC in responders than in proposers.

To identify edges showing significant role-related differences in ICC within each dataset, a permutation-based null model was constructed separately for NP-NR and NP-AR. Within each condition, proposer and responder labels were randomly shuffled across the pooled set of participants from that dataset while preserving the original group sizes. After each shuffle, edge-wise ICC was recomputed for the permuted proposer and responder sets, and a null difference map was obtained.

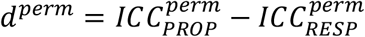

This procedure was repeated 1000 times for each condition. Directional empirical p-values were then derived separately for the two possible contrasts. Evidence for RESP > PROP effects was assessed from the negative tail of the observed difference map, whereas evidence for PROP > RESP effects was assessed from the positive tail. False discovery rate (FDR) correction was applied separately to the two directional p-value vectors, with a corrected threshold of q≤0.01

For visualisation and analyses, significant edges were encoded in sign masks. Edges with significantly higher ICC in responders than in proposers were assigned a value of +1 (RESP > PROP), whereas edges with significantly higher ICC in proposers than in responders were assigned a value of -1 (PROP > RESP). Furthermore, the spatial distribution of significant edges was summarised at the canonical network level by aggregating the significantly different edges into 7 × 7 matrices based on Yeo network assignments^36^.

### Network-level comparison of role-difference ICC between datasets

To test whether proposer versus responder ICC differences differed between the NP-NR and NP-AR datasets, an additional comparison was performed at the network-pair level using the pooled residualised data. This analysis was based on the dataset-specific role-difference maps

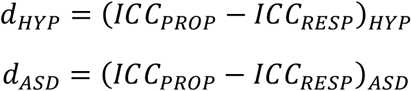

and on their between-dataset difference

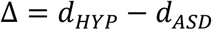

computed separately for each condition and for the condition-averaged map. The resulting edge-wise Δ maps were aggregated into 7 × 7 canonical network-pair matrices.

Statistical significance was assessed using a permutation procedure that shuffled NP-NR versus NP-AR labels within role, so that proposer labels were permuted only among proposers and responder labels only among responders. This preserved the proposer-responder structure while testing whether dataset membership contributed to the observed difference in role-related ICC. For each permutation, ICC was recomputed for the relabelled NP-NR and NP-AR datasets, the corresponding Δperm map was obtained, and its network-level 7 × 7 summary was stored. This procedure was repeated 1000 times. Empirical directional p-values were then computed for each network cell, followed by false discovery rate correction at q ≤ 0.01. Positive cells indicated NP-NR > NP-AR, whereas negative cells indicated NP-AR > NP-NR in the proposer-versus-responder ICC contrast.

### Edge-wise spatial specificity for autistic-traits analyses

For the analyses relating differential ICC patterns to autistic traits, edge-wise spatial specificity was also estimated in the NP-AR-only motion-residualised data. In this case, the same ICC stabilisation and permutation framework described above was applied to NP-AR proposers and NP-AR responders only. Thus, for each condition, edge-wise ICC was estimated separately in NP-AR proposers and NP-AR responders by repeated 80% subsampling across 100 iterations, and the role-difference map was computed

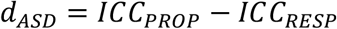

Permutation-based null distributions were generated by shuffling proposer and responder labels within the pooled NP-AR sample, separately for each condition, with 1000 permutations. Directional p-values were computed exactly as in the pooled analysis, and false discovery rate correction was applied separately for the responder-greater-than-proposer and proposer-greater-than-responder directions at q ≤ 0.01. Sign masks were again encoded such that +1 corresponded to RESP > PROP and −1 to PROP > RESP.

To further characterise the functional relevance of the spatially distributed differential ICC patterns, meta-analytic functional decoding was performed using the Neurosynth platform (https://neurosynth.org/). Following procedures established in previous work^20,22^, ROI inclusion masks were generated separately for the RESP > PROP and PROP > RESP directions in CTRL, PP, PR, and in the condition-averaged map. Each mask was then evaluated against a predefined set of 21 topic terms derived from the 50-topic Neurosynth model, selected to represent broad cognitive and functional domains. For each topic and mask, the corresponding Neurosynth decoding z-statistic was extracted and arranged into topic-by-condition summary heatmaps, separately for the two contrasts. Only decoding values meeting the predefined threshold of z ≥ 3.1 were retained.

In addition, a leave-one-out version of the NP-AR-only procedure was implemented. For each fold, one participant was excluded, edge-wise ICC was recomputed in the remaining training sample by the same bootstrap-subsampling procedure, and permutation-based sign masks were generated from the training data only. The left-out participant was never used in the computation of the corresponding mask. This produced fold-specific leave-one-out masks for CTRL, PP, PR, and their condition-averaged summary, which were then applied in the subsequent AQ analyses, including both linear regression and classification models to avoid data leakage.

### Analyses of autistic traits

Associations between autistic traits and differential ICC-derived connectivity features were examined in the NP-AR dataset. AQ scores were analysed both as a continuous outcome in regression models and, in a secondary analysis, as a binary label obtained by thresholding AQ scores at 26 for classification analyses.

The first set of analyses was based on the NP-AR-only motion-residualised data from Set B. For each condition and for the condition-averaged map, two common differential ICC masks were available, corresponding to edges with significantly greater ICC in responders than in proposers and to edges with significantly greater ICC in proposers than in responders. For each participant, FC matrices were first averaged across test and retest halves within condition. Subject-level masked FC features were then computed as the mean absolute FC magnitude across all edges belonging to a given mask. This produced, for each participant, one summary value for the responder-greater-than-proposer mask and one summary value for the proposer-greater-than-responder mask in each of CTRL, PP, PR, and the condition-averaged map. These masked FC summaries were then related to AQ scores using Pearson correlation, with FDR correction applied across the set of mask/condition tests.

The leave-one-out fold-specific Set B masks described in the previous section were first used in linear models predicting continuous AQ scores. For each fold, the masked predictor for the held-out participant was defined as the mean absolute FC value across the edges selected by that participant’s corresponding leave-one-out mask. A linear model was then fitted in the remaining participants and used to predict the AQ score of the held-out subject. This procedure was repeated for both mask directions and for CTRL, PP, PR, and the condition-averaged map, resulting in eight linear models. Predictive performance was quantified as the Pearson correlation between observed and predicted AQ scores, together with the corresponding coefficient of determination and mean absolute error (MAE). P-values were adjusted by FDR correction.

A further analysis examined whether AQ threshold status could be classified from topology features derived from leave-one-out condition-averaged differential ICC masks. This analysis was restricted to NP-AR responders and used the residualised data from Set C.

For each leave-one-out fold, topology features were extracted from responder-specific FC matrices separately for CTRL, PP, and PR using fold-specific condition-averaged differential ICC masks. Both RESP > PROP and PROP > RESP mask directions were evaluated.

Classification performance was therefore assessed across the different FC condition and mask-direction combinations using the same nested leave-one-out framework. The fold-specific condition-averaged leave-one-out masks, previously derived from the NP-AR-only proposer-versus-responder differential ICC procedure, were applied to the responder FC matrices. Topology features were extracted separately from the two directional masked graphs corresponding to RESP > PROP and PROP > RESP edges. The selected metrics were chosen to provide non-redundant coverage of three key topological domains: centrality, integration, and segregation, along with a measure of network organisation. Specifically, a representative measure was extracted for each domain: for centrality, eigenvector centrality; for integration, characteristic path length; for segregation, modularity. Additionally, network organisation was quantified using assortativity.

Classification analyses were then performed to predict whether AQ was greater than or equal to 26. Nested leave-one-out cross-validation was used, with mask generation and model fitting performed independently within each outer fold. A linear support vector machine (SVM) with class balancing was used as the classifier. Hyperparameters were tuned in an inner stratified cross-validation loop using the area under the receiver operating characteristic curve (ROC-AUC) as the optimisation criterion. Outer-loop performance was quantified using accuracy, balanced accuracy, sensitivity, specificity, F1-score, Matthews correlation coefficient (MCC), and AUC.

Additional validation analyses were performed around the classification framework. These included fold-wise export of SVM predictions, influence diagnostics based on removing one predicted subject at a time from the set of outer-loop predictions, univariate receiver operating characteristic analyses for each feature separately, and inspection of feature intercorrelations. Statistical significance of classification performance was assessed with a label permutation test (5000 permutations) that reran the full nested leave-one-out procedure under shuffled AQ labels and compared the observed outer-loop area under the receiver operating characteristic curve against the resulting empirical null distribution.

To assess the sensitivity of the best performing classification result to the choice of AQ threshold, the same classification framework was repeated using AQ thresholds from 23 to 29. This analysis was restricted to the best performing model identified in the primary analysis. For each distinct threshold-defined class assignment, the complete nested leave-one-out procedure and the 5000 permutations significance test were repeated.

