## Supplementary Materials for "Social brain fingerprinting: unique functional connectomes underpin interpersonal behaviours in non-autistic and autistic adults"

[1] IRCCS Centro Neurolesi "Bonino Pulejo", 98124 Messina, Italy.

[2] Behavioural and Social Neuroscience Research Group, Central European Institute of Technology (CEITEC), Masaryk University, Brno, Czechia

[3] Institute of Psychology, Czech Academy of Sciences, Brno, Czechia

[4] Brain Mapping Lab, Department of Biomedical, Dental Sciences and Morphological and Functional Imaging, University of Messina, Messina, Italy

[5] Department of Biomedical Sciences, Humanitas University, Via Rita Levi Montalcini 4, Pieve Emanuele, 20072 Milan, Italy

[6] IRCCS Humanitas Research Hospital, Via Alessandro Manzoni 56, Rozzano, 20089 Milan, Italy

[7] First Department of Neurology, St. Anne's University Hospital and Faculty of Medicine,

Masaryk University, Brno, Czechia

[8] Department of Psychology, Aston University, Birmingham, UK

[9] Institute of Health and Neurodevelopment, Aston University, Birmingham, UK

[10] School of Mathematics, University of Birmingham, Birmingham, UK

[11] Centre for Human Brain Health, University of Birmingham, Birmingham, UK

[§] Shared co-last authorship

[*] Correspondence

**Augusto Ielo**

IRCCS Centro Neurolesi "Bonino Pulejo", 98124 Messina, Italy.

**Enrico Amico**

School of Mathematics & Centre for Human Brain Health

University of Birmingham, B15 2TT Birmingham, UK

### Supplementary Materials

#### Table S1. Across-condition comparisons of whole-brain identifiability within group

| Dataset | Role | Contrast | Metric | Mean 1 | Mean 2 | Diff12 | t | d.f. | p, paired | p, permutation | q, FDR |
|---|---|---|---|---|---|---|---|---|---|---|---|
| NP-NR | PROP | PP VS CTRL | Iself | 0.59 | 0.54 | 0.05 | 2.43 | 18 | 0.026 | 0.025 | 0.311 |
| NP-NR | PROP | PP VS CTRL | Iothers | -0.01 | -0.01 | 0.00 | -1.03 | 18 | 0.315 | 0.317 | 0.630 |
| NP-NR | PROP | PR VS CTRL | Iself | 0.59 | 0.54 | 0.05 | 2.00 | 18 | 0.061 | 0.059 | 0.368 |
| NP-NR | PROP | PR VS CTRL | Iothers | -0.02 | -0.01 | -0.01 | -1.31 | 18 | 0.206 | 0.211 | 0.551 |
| NP-NR | PROP | PP vs PR | Iself | 0.59 | 0.59 | 0.00 | -0.35 | 18 | 0.733 | 0.735 | 0.799 |
| NP-NR | PROP | PP vs PR | Iothers | -0.02 | -0.01 | -0.01 | -0.37 | 18 | 0.715 | 0.711 | 0.858 |
| NP-NR | RESP | PP VS CTRL | Iself | 0.58 | 0.56 | 0.02 | 0.77 | 18 | 0.454 | 0.442 | 0.663 |
| NP-NR | RESP | PP VS CTRL | Iothers | -0.01 | -0.01 | 0.00 | 0.03 | 18 | 0.973 | 0.971 | 0.973 |
| NP-NR | RESP | PR VS CTRL | Iself | 0.55 | 0.56 | -0.01 | -0.61 | 18 | 0.553 | 0.555 | 0.663 |
| NP-NR | RESP | PR VS CTRL | Iothers | 0.00 | -0.01 | 0.01 | 4.61 | 18 | <0.001 | <0.001 | 0.001 |
| NP-NR | RESP | PP vs PR | Iself | 0.55 | 0.58 | -0.03 | -1.31 | 18 | 0.205 | 0.205 | 0.462 |
| NP-NR | RESP | PP vs PR | Iothers | 0.00 | -0.01 | 0.01 | 4.74 | 18 | <0.001 | <0.001 | 0.001 |
| NP-AR | PROP | PP VS CTRL | Iself | 0.59 | 0.56 | 0.03 | 1.03 | 15 | 0.320 | 0.318 | 0.549 |
| NP-AR | PROP | PP VS CTRL | Iothers | -0.02 | -0.01 | -0.01 | -0.58 | 15 | 0.569 | 0.557 | 0.781 |
| NP-AR | PROP | PR VS CTRL | Iself | 0.59 | 0.56 | 0.03 | 1.25 | 15 | 0.231 | 0.228 | 0.462 |

| | | | | | | | | | | | |
|---|---|---|---|---|---|---|---|---|---|---|---|
| NP-AR | PROP | PR VS CTRL | Iothers | -0.01 | -0.01 | 0.00 | -0.64 | 15 | 0.534 | 0.529 | 0.781 |
| NP-AR | PROP | PP vs PR | Iself | 0.59 | 0.59 | 0.00 | 0.12 | 15 | 0.903 | 0.900 | 0.903 |
| NP-AR | PROP | PP vs PR | Iothers | -0.01 | -0.02 | 0.01 | 0.08 | 15 | 0.941 | 0.936 | 0.973 |
| NP-AR | RESP | PP VS CTRL | Iself | 0.60 | 0.58 | 0.02 | 1.31 | 15 | 0.209 | 0.199 | 0.462 |
| NP-AR | RESP | PP VS CTRL | Iothers | -0.01 | -0.01 | 0.00 | 0.56 | 15 | 0.586 | 0.582 | 0.781 |
| NP-AR | RESP | PR VS CTRL | Iself | 0.56 | 0.58 | -0.02 | -0.67 | 15 | 0.510 | 0.518 | 0.663 |
| NP-AR | RESP | PR VS CTRL | Iothers | -0.00 | -0.01 | 0.01 | 1.76 | 15 | 0.098 | 0.097 | 0.393 |
| NP-AR | RESP | PP vs PR | Iself | 0.56 | 0.60 | -0.04 | -1.61 | 15 | 0.128 | 0.130 | 0.462 |
| NP-AR | RESP | PP vs PR | Iothers | -0.00 | -0.01 | 0.01 | 1.25 | 15 | 0.230 | 0.224 | 0.551 |

*Legend: Across-condition comparisons were performed within each group using two-sided paired t-tests and sign-flip permutation tests. Mean 1 and Mean 2 correspond to the first and second conditions listed in the contrast, respectively, and Diff12 represents Mean 1 − Mean 2. q values represent false discovery rate-adjusted p-values. False discovery rate correction was applied separately within the Iself and Iothers families. Iself, self-identifiability; Iothers, others-identifiability.*

**Table S2. Between-dataset comparisons of whole-brain identifiability within role**

| Contrast | Role | Condition | Metric | Mean 1 | Mean 2 | Diff12 | t | d.f. | p, Welch | p, permutation | q, FDR |
|---|---|---|---|---|---|---|---|---|---|---|---|
| NP-NR vs NP-AR | PROP | CTRL | Iself | 0.54 | 0.56 | -0.02 | -0.58 | 25.31 | 0.567 | 0.559 | 0.968 |
| NP-NR vs NP-AR | PROP | CTRL | Iothers | -0.01 | -0.01 | 0.00 | 0.11 | 25.15 | 0.917 | 0.916 | 0.963 |
| NP-NR vs NP-AR | RESP | CTRL | Iself | 0.56 | 0.58 | -0.02 | -0.63 | 32.72 | 0.531 | 0.541 | 0.968 |
| NP-NR vs NP-AR | RESP | CTRL | Iothers | -0.01 | -0.01 | 0.00 | 0.36 | 28.99 | 0.720 | 0.718 | 0.963 |
| NP-NR vs NP-AR | PROP | PP | Iself | 0.59 | 0.59 | 0.00 | 0.21 | 22.33 | 0.833 | 0.832 | 0.968 |
| NP-NR vs NP-AR | PROP | PP | Iothers | -0.01 | -0.02 | 0.01 | 0.05 | 24.02 | 0.963 | 0.964 | 0.963 |
| NP-NR vs NP-AR | RESP | PP | Iself | 0.58 | 0.60 | -0.02 | -0.74 | 31.93 | 0.463 | 0.475 | 0.968 |
| NP-NR vs NP-AR | RESP | PP | Iothers | -0.01 | -0.01 | 0.00 | 0.17 | 28.95 | 0.863 | 0.862 | 0.963 |
| NP-NR vs NP-AR | PROP | PR | Iself | 0.59 | 0.59 | 0.00 | -0.04 | 32.99 | 0.968 | 0.971 | 0.968 |
| NP-NR vs NP-AR | PROP | PR | Iothers | -0.02 | -0.01 | -0.01 | -0.08 | 22.95 | 0.936 | 0.934 | 0.963 |
| NP-NR vs NP-AR | RESP | PR | Iself | 0.55 | 0.56 | -0.01 | -0.33 | 29.08 | 0.744 | 0.746 | 0.968 |
| NP-NR vs NP-AR | RESP | PR | Iothers | 0.00 | 0.00 | 0.00 | 0.65 | 26.26 | 0.521 | 0.499 | 0.963 |

*Legend: Between-dataset comparisons were performed within role using two-sided Welch two-sample t-tests and label-shuffling permutation tests. Mean 1 and Mean 2 correspond to NP-NR and NP-AR, respectively, and Diff12 represents Mean 1 − Mean 2. q values represent false discovery rate-adjusted p-values. False discovery rate correction was applied separately within the Iself and Iothers families. Iself, self-identifiability; Iothers, others-identifiability.*

**Table S3. Sensitivity of AQ-status classification to alternative AQ thresholds**

| AQ threshold | Lower AQ, n | Higher AQ, n | Changed vs AQ≥26, n | Accuracy | Balanced accuracy | AUC | Sensitivity | Specificity | F1 | MCC | p, permutation (AUC) |
|---|---|---|---|---|---|---|---|---|---|---|---|
| ≥23 | 5 | 10 | 1 | 0.87 | 0.90 | 0.88 | 0.80 | 1.00 | 0.89 | 0.76 | 0.012 |
| ≥24 | 5 | 10 | 1 | 0.87 | 0.90 | 0.88 | 0.80 | 1.00 | 0.89 | 0.76 | 0.012 |
| ≥25 | 6 | 9 | 0 | 0.87 | 0.89 | 0.85 | 0.78 | 1.00 | 0.88 | 0.76 | 0.021 |
| ≥26 | 6 | 9 | 0 | 0.87 | 0.89 | 0.85 | 0.78 | 1.00 | 0.88 | 0.76 | 0.021 |
| ≥27 | 8 | 7 | 2 | 0.53 | 0.53 | 0.63 | 0.43 | 0.63 | 0.46 | 0.05 | 0.243 |
| ≥28 | 8 | 7 | 2 | 0.53 | 0.53 | 0.63 | 0.43 | 0.63 | 0.46 | 0.05 | 0.243 |
| ≥29 | 9 | 6 | 3 | 0.67 | 0.67 | 0.78 | 0.67 | 0.67 | 0.62 | 0.33 | 0.066 |

*Legend: Performance of the best-performing linear support vector machine across alternative Autism-Spectrum Quotient (AQ) cutoffs. The model used PR functional-connectivity topology features derived from the condition-averaged RESP > PROP differential ICC mask. Classification was evaluated using nested leave-one-out cross-validation. Statistical significance of outer-loop ROC-AUC was assessed using 5000 label permutations in which the complete nested cross-validation procedure, including inner-loop hyperparameter optimisation, was repeated. "Changed vs AQ≥26" indicates the number of autistic Responders whose binary class assignment differed from the primary AQ≥26 definition. Because no participants had AQ values separating some adjacent thresholds, AQ≥23 and ≥24, AQ≥25 and ≥26, and AQ≥27 and ≥28 generated identical subject-level labels and therefore identical classification results; equivalent models were evaluated only once. The AQ≥26 row corresponds to the primary classification analysis reported in the main text. Permutation p-values refer to the one-sided empirical test of the outer-loop AUC against the permutation-derived null distribution.*

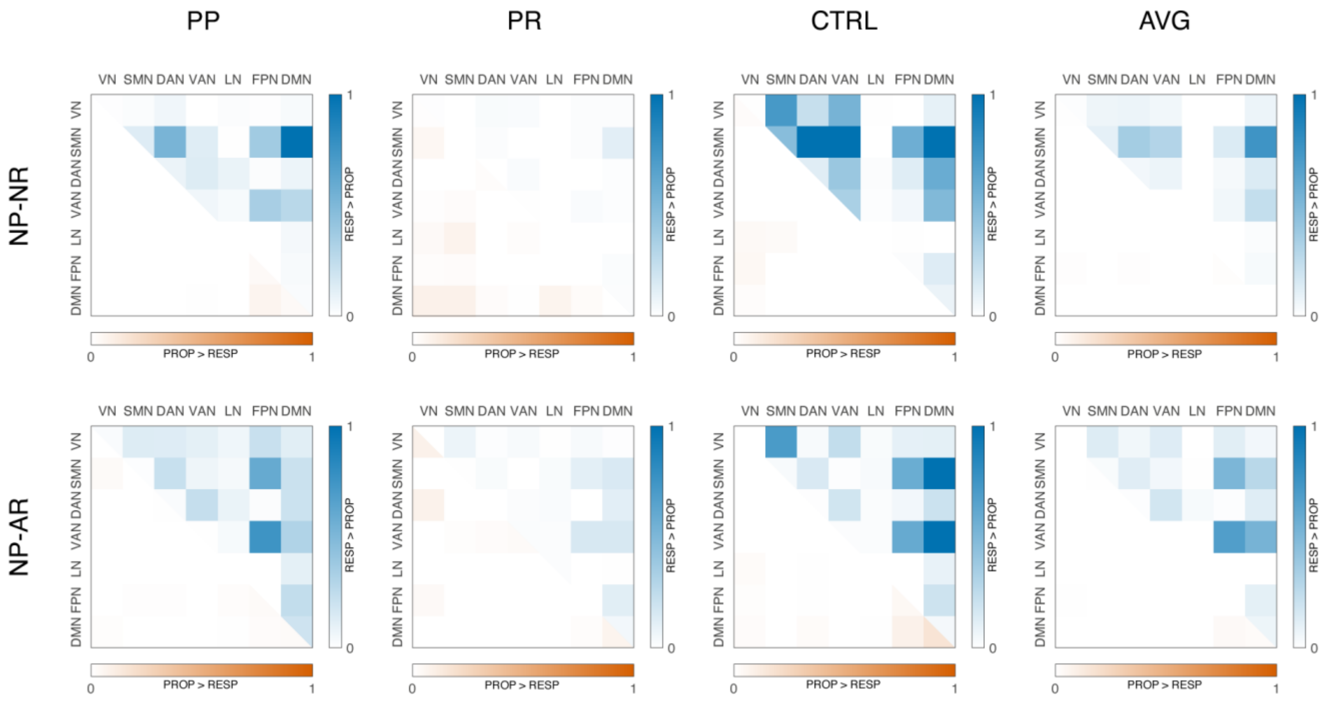


**Figure S1. Network-level summaries of within-dataset proposer–responder ICC differences.** Each panel shows the network-level distribution of significant edge-wise ICC differences for CTRL, PP, PR and the condition-averaged map (AVG), separately for NP-NR and NP-AR. Blue indicates network pairs in which responders showed higher ICC than proposers (RESP > PROP), whereas orange indicates network pairs in which proposers showed higher ICC than responders (PROP > RESP). Colour intensity reflects the normalised magnitude of the corresponding network-level effect.

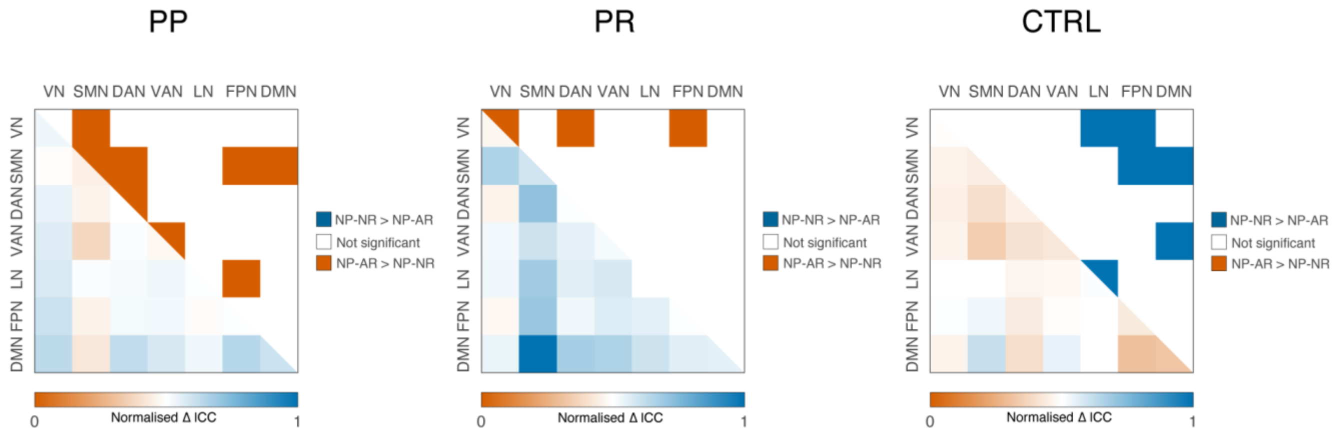


**Figure S2. Network-level comparison of proposer–responder ICC differences between NP-NR and NP-AR.** Each panel shows the between-dataset difference in the role-difference ICC map for one condition (PP, PR, CTRL). In each matrix, the lower triangle displays the normalised network-level ΔICC, where blue indicates larger proposer–responder ICC differences in NP-NR than in NP-AR, and orange indicates larger proposer–responder ICC differences in NP-AR than in NP-NR. Colour intensity in the lower triangle reflects the normalised magnitude of the between-dataset effect on a common 0–1 scale across conditions. The upper triangle displays the corresponding FDR-corrected significance mask from the permutation test: blue = significant NP-NR > NP-AR, orange = significant NP-AR > NP-NR, white = not significant. Here, ΔICC denotes the between-dataset difference in the proposer–responder ICC contrast.

**List of abbreviations**

AQ, Autism-Spectrum Quotient; AUC, area under the receiver operating characteristic curve; AVG, condition-averaged map; CTRL, control condition; DAN, dorsal attention network; d.f., degrees of freedom; DMN, default mode network; F1, F1-score; FDR, false discovery rate; FPN, frontoparietal network; ICC, intraclass correlation coefficient; Iothers, others-identifiability; Iself, self-identifiability; LN, limbic network; MCC, Matthews correlation coefficient; NP-AR, non-autistic Proposer–autistic Responder dataset; NP-NR, non-autistic Proposer–non-autistic Responder dataset; PP, Proposer–Proposer condition; PR, Proposer–Responder condition; PROP, Proposer; RESP, Responder; ROC-AUC, receiver operating characteristic area under the curve; SMN, somatomotor network; VAN, ventral attention network; VN, visual network.
